# Single-subject Proteomic Signatures in Alzheimer’s Disease Reflect Clinical Phenotypes and Distinguish Asymptomatic from Symptomatic Cases

**DOI:** 10.64898/2026.02.06.704408

**Authors:** Avijit Podder, Yi Juin Liew, Greg A. Cary, Gregory W. Carter, Asli Uyar

## Abstract

**INTRODUCTION:** Alzheimer’s Disease (AD) exhibits considerable inter-individual variability in clinical presentation, neuropathological burden, and underlying molecular processes. Conventional cohort-based analyses of omics molecular data often mask individual-level heterogeneity, limiting insights into the precision therapeutic strategies. To address this challenge, we developed INDIGO (INdividual-level DIfferential GenOmics), a computational framework that quantifies molecular deviations for each individual relative to healthy controls, enabling subject-specific profiling of disease-associated alterations in proteomic data, with a framework that is readily applicable to other omics modalities.

**METHODS:** We applied INDIGO to dorsolateral prefrontal cortex (DLPFC) proteomic data from ROSMAP cohort (N = 610). Protein level deviations were aggregated into gene set activity scores for KEGG pathways and curated AD Biodomain annotations. Functional alterations across AD and Asymptomatic AD (AsymAD) individuals were evaluated and correlated with clinical metrics including APOE4 genotype, Braak stage, CERAD and MMSE scores. Graph-based clustering was used to identify molecularly distinct subgroups based on shared patterns of functional dysregulation.

**RESULTS:** Limited overlap was observed between cohort-level differential expression analysis and INDIGO single-subject analyses. Individual deviations in various processes, including metabolic, immune and epigenetic pathways, exhibited sex- and disease stage-specific patterns. Amyloid clearance and immune activation were strongly associated with APOE4 dosage, higher amyloid and tau burden and cognitive decline, whereas upregulation of mitochondrial and synaptic modules correlated positively with preserved cognitive function. By linking individuals through concordant directional proteomic changes, we identified molecularly coherent subgroups that transcend conventional diagnostic boundaries and include both AD and AsymAD subjects. Each subgroup displayed distinct functional signatures and defined by a unique set of key regulatory proteins.

**DISCUSSION:** These results demonstrate that single-subject omics profiling can resolve individual molecular signatures aligned with clinical and neuropathological variation in AD. By linking molecular heterogeneity with disease phenotypes, INDIGO provides a scalable framework for precision modeling and novel therapeutic target discovery.

## 1. BACKGROUND

Alzheimer’s disease (AD) is a progressive neurodegenerative disorder and the leading cause of dementia worldwide. Its clinical manifestations are highly heterogeneous with variation in the age of onset, early symptoms, neuropathological characteristics and rates of cognitive decline, following distinct disease progression trajectories^1-3^.This variability complicates prognosis and treatment, highlighting the need to dissect molecular mechanisms driving divergent disease outcomes.

Recent advances in omics technologies enabled probing AD heterogeneity at molecular, cellular and network levels and uncovering molecular subtypes that align with clinical phenotypes^4^. Transcriptomic and multi-omic clustering approaches revealed subgroups of AD patients characterized by divergent molecular signatures recapitulating known pathological features and highlight key processes like lipid metabolism, mitochondrial decline, and epigenetic regulation that contribute to disease variability.^5-7^

While group-level subtyping has deepened our understanding of AD heterogeneity, majority of these strategies are inherently top-down: they cluster patients by partitioning omics data across the study population and then identify biological features that distinguish groups. Such population-average approaches often overlook inter-individual variation that shapes disease trajectories. Alternative methodologies, such as individualized pathway aberrance score (iPAS)^8^, ‘N-of-1’ pathways^9^ and personalized perturbation profiles (PEEPs)^10^, address this gap by computing gene or pathway level molecular deviations for each individual and enable direct linkage of omics signatures to clinical phenotypes^11^.

Among omics layers, proteomics offers the most functionally proximal readout of cellular state capturing post-transcriptional regulation and protein-level changes that more directly reflect disease mechanisms and therapeutic targets. Proteomic profiling is especially relevant in neurodegenerative disorders, where protein aggregation, synaptic dysfunction, and inflammation are driven by alterations beyond the transcriptome^12, 13^. Although most single-subject approaches were originally developed using transcriptomic data, they are adaptable to diverse omics and multi-omics profiles^11^ and align closely with precision medicine objectives, however, their utilization in AD remains limited.

To provide a more precise characterization of AD heterogeneity, here we introduce INDIGO (INdividual-level DIfferential GenOmics), a computational framework to quantify molecular deviations in individual AD samples relative to cognitively unimpaired control group. We applied INDIGO to dorsolateral prefrontal cortex (DLPFC) proteomes from the Religious Orders Study and Memory and Aging Project (ROSMAP) cohort^14^. The initial step, single-subject differential expression analysis (ssDEA), derives protein-level deviation profiles, capturing how each individual’s molecular state diverges from reference patterns and transforms raw proteomes into individualized molecular signatures^10^. Subsequently, single-subject functional profiling analysis (ssFPA), integrates these deviations across curated biological pathways and functional gene sets (e.g., KEGG, GO Biological Process and AD Biodomains^15^). This generates a subject-by-pathway activity landscape, enabling the identification of dysregulated biological processes underlying disease heterogeneity.

The resulting individual-level molecular signatures distinguish asymptomatic from symptomatic AD, reveal sex-stratified pathways, correlate with clinical metrics and identify subgroups defined by co-regulated biological processes. These findings illustrate the potential of single-subject proteomics to bridge molecular heterogeneity with clinical phenotypes and advance precision medicine in AD.

## 2. METHODS

### 2.1 Data collection

We analyzed tandem mass tag (TMT)–based proteomic data generated from postmortem DLPFC (Brodmann area 9) tissue of 610 participants enrolled in the ROSMAP study, obtained from the AD Knowledge Portal (ADKP) (Synapse ID: syn31534849). Clinical and neuropathological annotations followed established ROSMAP protocols, including Mini-Mental State Examination (MMSE), CERAD neuritic plaque score, and Braak neurofibrillary staging^16^.

AD cases were defined by dementia (MMSE < 24) with moderate-to-severe neuropathology (Braak stage 3–6; CERAD 1–2). Asymptomatic AD (AsymAD) individuals met similar neuropathological criteria but were cognitively unimpaired (MMSE ≥ 24). Control subjects showed minimal pathology (Braak stage 0–3; CERAD 3–4) and intact cognition. Samples not meeting these criteria were excluded. This classification scheme is consistent with prior large-scale ROSMAP proteomic studies^13^ and is visually summarized in Figure 1A. Final sample group distributions and counts are presented in Figure 1B.

**Figure 1.**
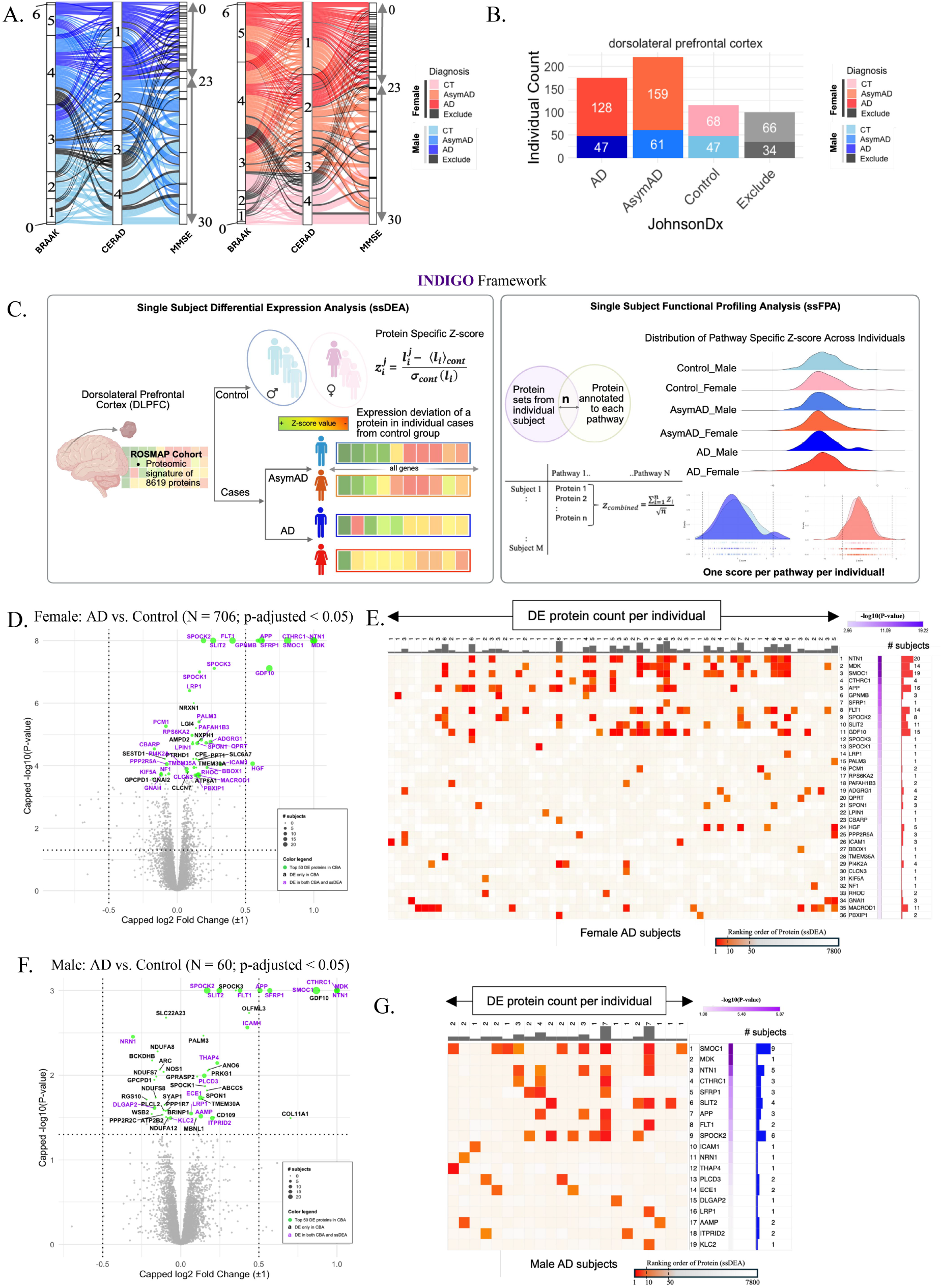
INDIGO framework, study population, and comparison of cohort-level vs. single-subject differential expression. **(A)** Schematic of the INDIGO workflow: protein-level z-scores are computed for each subject relative to cognitively normal controls, followed by single-subject functional pathway analysis (ssFPA). **(B)** Sankey diagram illustrating the classification of ROSMAP participants into AD, AsymAD, and Control groups, integrating neuropathological and cognitive criteria (per ^13^). **(C)** Distribution of proteomic samples by diagnosis and sex, showing balanced representation that enables sex-stratified single-subject profiling. **(D, F)** Volcano plots of cohort-level differential expression (DE) for AD versus controls in females (D) and males (F). Proteins in purple indicate overlap with the top 50 most deviating proteins in at least one AD subject identified by INDIGO. **(E, G)** Heatmaps of within-subject ranks for the top 50 DE proteins in female (E) and male (G) AD subjects who exhibited at least one of these proteins among their individual top 50 deviating proteins. Individuals (columns) were ordered using hierarchical clustering of protein z-scores. Most proteins highlighted at the cohort level were not consistently prioritized across individuals, underscoring substantial inter-individual heterogeneity in proteomic alterations.

Protein abundance data were obtained following standardized ROSMAP preprocessing, including pooled-reference normalization, missingness filtering, and batch correction. After quality control and redundancy resolution, the final dataset comprised approximately 7,000–8,000 proteins per individual and was used for all downstream analyses (Supplementary Methods 2.1 and Figures S1).

### 2.2 Single-subject Differential Expression Analysis (ssDEA)

To quantify individual-level proteomic deviations, we performed single-subject differential expression analysis (ssDEA) using protein-wise z-score calculations. Analyses were conducted on log□-transformed, TAMPOR-corrected protein abundance data obtained from the ADKP. For each AD and AsymAD individual, protein z-scores were computed relative to sex-matched cognitively normal controls, enabling standardized assessment of subject-specific molecular deviations^10^. The overall analysis workflow for ssDEA is illustrated in Figure 1C.

For a given protein *i* in subject *j*, the z-score was calculated as:

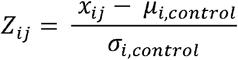

where *x*_*ij*_ denotes the abundance of protein *i* in subject *j*, and *µ*_*i,control*_, *σ*_*i,control*_ represent the mean and standard deviation of that protein among sex-matched control subjects.

To ensure a robust reference baseline, internal variability within the control cohort was evaluated by leave-one-out z-score analysis (Supplementary Methods 2.2 and Figures S2–S3), with missing values left un-imputed to avoid bias, consistent with proteomics best practices.^17^

### 2.3 Comparison with Cohort-Based Differential Expression Analysis

To evaluate how ssDEA differs from conventional population-level approaches, we compared ssDEA results with cohort-based DE analyses of AD versus control subjects. Sex-stratified cohort-level DE results were obtained from the ADKP (Synapse ID: syn25607662).^13^ For each sex, top 50 proteins identified by adjusted p-values in the cohort-based analysis were compared with the top 50 proteins ranked by ssDEA z-scores at the individual level. The degree of overlap between these lists was used to quantify concordance between population-derived and individualized proteomic signatures.

### 2.4 Single-subject Functional Profiling Analysis (ssFPA)

To characterize subject-specific functional alterations, we implemented ssFPA by integrating individual protein z-scores with curated functional annotations (Supplementary Methods 2.4). Two complementary annotation frameworks were used: (i) 347 canonical pathways from KEGG, and (ii) an AD-specific functional framework comprising 18 high-level biodomains^15^ and 107 nested subdomains (Synapse ID: syn26529354) that capture molecular processes relevant to AD pathogenesis.

#### Quantification of Individual-Level Functional Perturbations

Pathway- and subdomain-level z-scores were computed for all individuals based on single-subject protein profiles. For each module, empirical reference bounds were defined using control subjects as the 1st–99th percentile range of z-scores. AD and AsymAD individuals with values above or below these bounds were classified as upregulation or downregulation, respectively. The proportion of individuals showing deregulation was quantified for each diagnostic group.

### 2.5 Association of Subdomain-Level Functional Dysregulation with Clinical and Neuropathological Metrics

To assess clinical relevance, we tested associations between subdomain-level z-scores and key clinical and neuropathological measures across all subjects. Analyses focused on APOE4 allele count, Braak stage, CERAD neuritic plaque score, and MMSE. Aggregating control, AsymAD and AD subjects for clinical correlations provided a continuum of disease progression, where AsymAD represents an intermediate or prodromal state^18, 19^, thereby increasing sensitivity to early or subtle disruptions.

Associations with ordinal traits were evaluated using nonparametric tests, while linear correlations were used for MMSE. Subdomains showing statistically significant associations after multiple-testing correction were considered clinically or pathologically relevant.

### 2.6 Graph-Based Clustering of Functional Signatures

To identify molecularly defined subgroups among AD and AsymAD subjects, we constructed a bipartite graph linking individuals to functionally dysregulated subdomains. For each subject, subdomains with z-scores outside the control-derived 1st–99th percentile range were selected as significantly perturbed. Subjects and subdomains were represented as nodes, with edges indicating significant dysregulation.

Community structure within this graph was identified using the Infomap algorithm, a flow-based community detection method that partitions networks based on information compression.^20^ This approach enables detection of molecularly coherent subgroups defined by shared functional dysregulation patterns across individuals (Supplementary Methods 2.7).

### 2.7 Clinical and Neuropathological Characterization of Functional Clusters

To assess the clinical and pathological relevance of graph-derived functional clusters, we compared demographic, clinical, and neuropathological measures across clusters, including age at death, APOE4 carrier status, Braak stage, CERAD score, postmortem interval, and MMSE. Pairwise differences were assessed using Wilcoxon rank-sum tests, with significance determined after multiple-testing correction.

### 2.8 Subdomain–Specific Protein–Protein Interaction Network Analysis

To contextualize subdomain-associated proteins within a functional interactome, we constructed brain-specific protein–protein interaction networks (PPIN) using experimentally validated human interactions.^21^ Subdomain-associated proteins were mapped onto this network, and topological analysis was used to identify highly connected hub proteins. Protein-level z-scores were summarized within diagnostic or cluster-defined groups and integrated with network connectivity to prioritize candidate regulators underlying subdomain-level functional dysregulation (Supplementary Methods 2.9).

### 2.9 Computational Framework and Data Visualization

All analyses were performed in R using reproducible workflows. Proteomic and clinical data were accessed from the ADKP, and custom scripts were used for single-subject z-score computation, functional aggregation, and downstream analyses. Graph-based clustering, network analyses and visualizations were conducted using standard R packages.

Further methodological details, parameter settings, and extended description of computational procedures are provided in the Supplementary Methods.

## 3 RESULTS

### 3.1 Limited overlap between group-based and individual-level proteomic signatures

We compared INDIGO-derived single-subject deviations with conventional cohort-based differential expression (DE) analyses between AD and control subjects (Figure 1D–G). While several proteins were consistently dysregulated across individuals (e.g., NTN1, MDK, SMOC1), substantial inter-individual heterogeneity was observed. Only ∼50% of AD subjects showed enrichment of cohort-level DE proteins among their top-ranked individual deviations, with similar patterns in males and females. Notably, several well-established AD-associated proteins, including LRP1, ICAM1 and FLT1^22-24^, were infrequently ranked among the most deregulated proteins at the individual level, highlighting variability in their contribution across subjects. This limited concordance persisted when extending the analysis beyond the top-ranked proteins (Figures S4–S5).

Together, these results demonstrate that cohort-level-averaged DE analyses mask substantial subject-specific proteomic variation. INDIGO captures this hidden heterogeneity, revealing individualized molecular signatures that are not apparent from cohort-level comparisons.

### 3.2 Differential pathway dysregulation across sex and disease stage

INDIGO pathway-level z-scores were used to quantify the prevalence of KEGG pathway dysregulation in AsymAD and AD relative to controls. Pathways altered in ≥10% of individuals revealed strong sex-stratified patterns, with additional differences between AsymAD and AD within each sex (Figure 2A). In males, pathway dysregulation was dominated by downregulation of metabolic processes, whereas females showed more prominent alterations in signaling-related pathways. Upregulated pathways in males were enriched for RNA processing and DNA repair, while females exhibited stronger dysregulation of pathways related to Wnt signaling, NF-κB signaling, and synaptic function.

**Figure 2.**
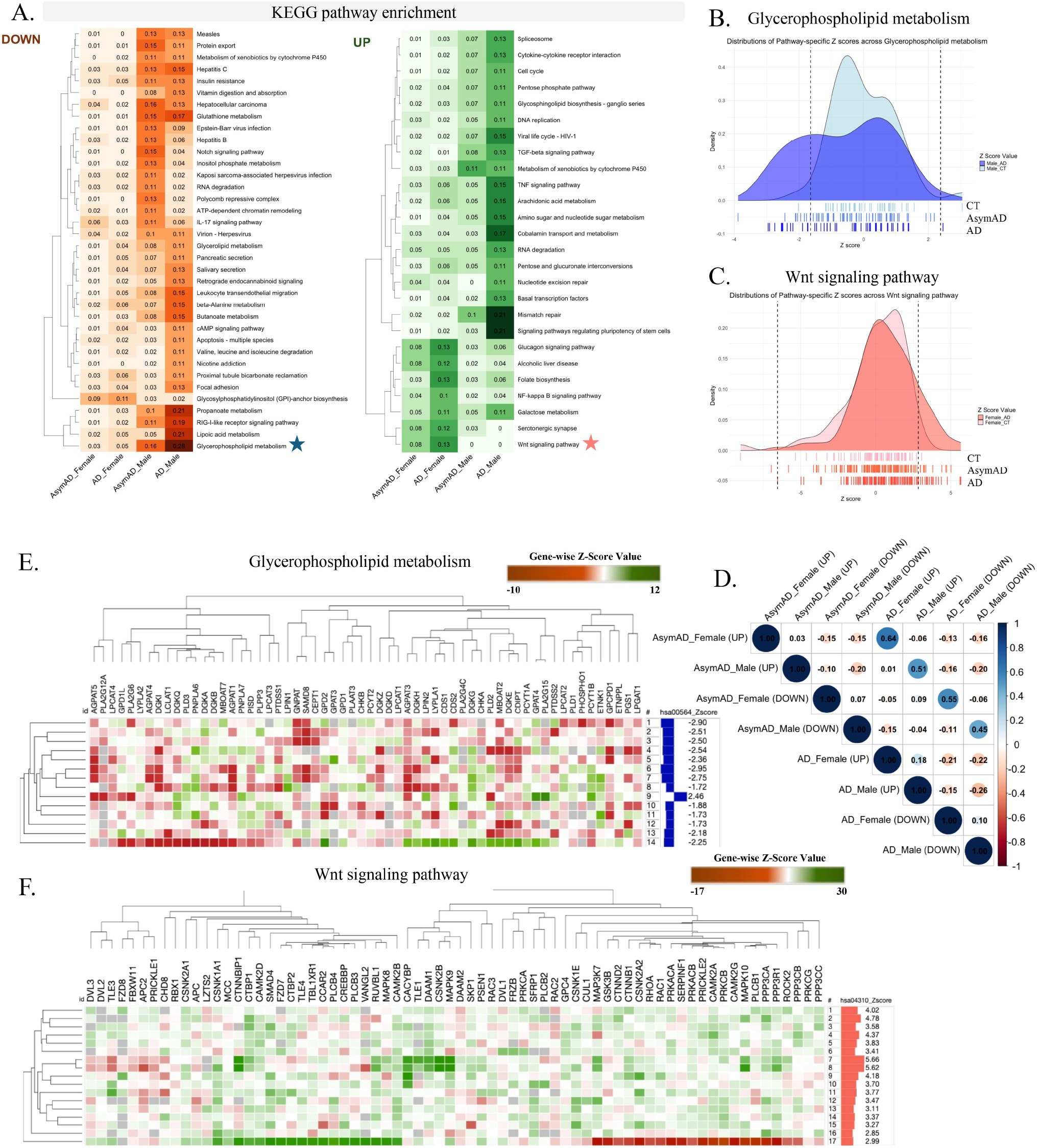
Pathway dysregulations across sex and disease stage. **(A)** Frequency of KEGG pathways showing significant single-subject deviations (beyond the control-derived 1st–99th percentile range) in AD and AsymAD individuals, stratified by sex. The enrichment patterns demonstrate a strong sex-separation and some differences between AD and AsymAD within the same sex. Downregulation was more prominent in males, glycerophospholipid, lipoic acid and propanoate metabolism being the top three pathways significant in >20% of male AD subjects and <=5% in female AD subjects. Only glycosylphosphatidylinositol (GPI)-anchor biosynthesis pathway was more frequently down-regulated in female AD (11%) compared to male AD (2%). Up-regulated pathways in males clustered around RNA processing and DNA repair (spliceosome, RNA degradation, mismatch repair, nucleotide excision repair). By contrast, females displayed stronger dysregulation of Wnt signaling, NF-κB, and serotonergic synapse pathways. **(B–C)** Density plots and individual z-score bars illustrating z-score distributions across groups for representative pathways: glycerophospholipid metabolism (B), predominantly downregulated in males, and Wnt signaling (C), predominantly upregulated in females. **(D)** Correlation of pathway enrichment frequencies between sex and diagnosis groups, highlighting shared alterations between AD and AsymAD within the same sex and divergent alterations between sexes. **(E–F)** Protein-level z-score heatmaps within exemplar pathways, showing heterogeneity at the protein level but pathway-consistent deviations: glycerophospholipid metabolism in males (E) and Wnt signaling in females (F). glycerophospholipid metabolism dysregulation in males was distributed across multiple proteins, supporting a pathway-level effect rather than a single-protein driver. Top downregulated proteins within this pathway include GK, DGKI and AGPAT1. Wnt signaling upregulation in females with heterogenous deviation patterns. Relatively few proteins including CTBP1, CAMK2B, CACYBP and DAAM1 were upregulated across individuals more consistently.

Two exemplar pathways illustrate these sex- and stage-specific trends. Glycerophospholipid metabolism was frequently downregulated in AD males, with AsymAD males showing intermediate profiles (Figure 2B). In contrast, Wnt signaling was prominently upregulated in AD females, partially altered in AsymAD females, and largely unchanged in males (Figure 2C). Although pathway enrichment frequencies were moderately correlated between AD and AsymAD groups, these relationships diverged when stratified by sex (Figure 2D).

Protein-level analyses within these pathways indicated that dysregulation reflected coordinated shifts across multiple proteins rather than dominance by single drivers (Figure 2E–F). Together, these findings demonstrate pronounced sex- and stage-dependent functional alterations in AD, with metabolic suppression predominating in males and aberrant signaling activation more prominent in females.

### 3.3 AD Biodomains distinguish asymptomatic and symptomatic AD

To examine higher-order functional organization, we applied INDIGO within the AD Biodomain framework, which groups AD-relevant processes into hierarchically structured biodomains and subdomains. Subdomain-level z-scores captured individual functional deviations across the disease spectrum (Figure S6).

Comparison of subdomain enrichment between AsymAD and AD subjects revealed distinct patterns. While amyloid-β formation was consistently altered in both groups, epigenetic subdomains showed opposing regulation, with suppression in AsymAD and activation in AD (Figure 3A). Transcription factor binding, histone modification, and miRNA-mediated silencing contributed most strongly to this divergence (Figure 3B), suggesting epigenetic remodeling as a key molecular feature distinguishing asymptomatic from symptomatic disease.

**Figure 3.**
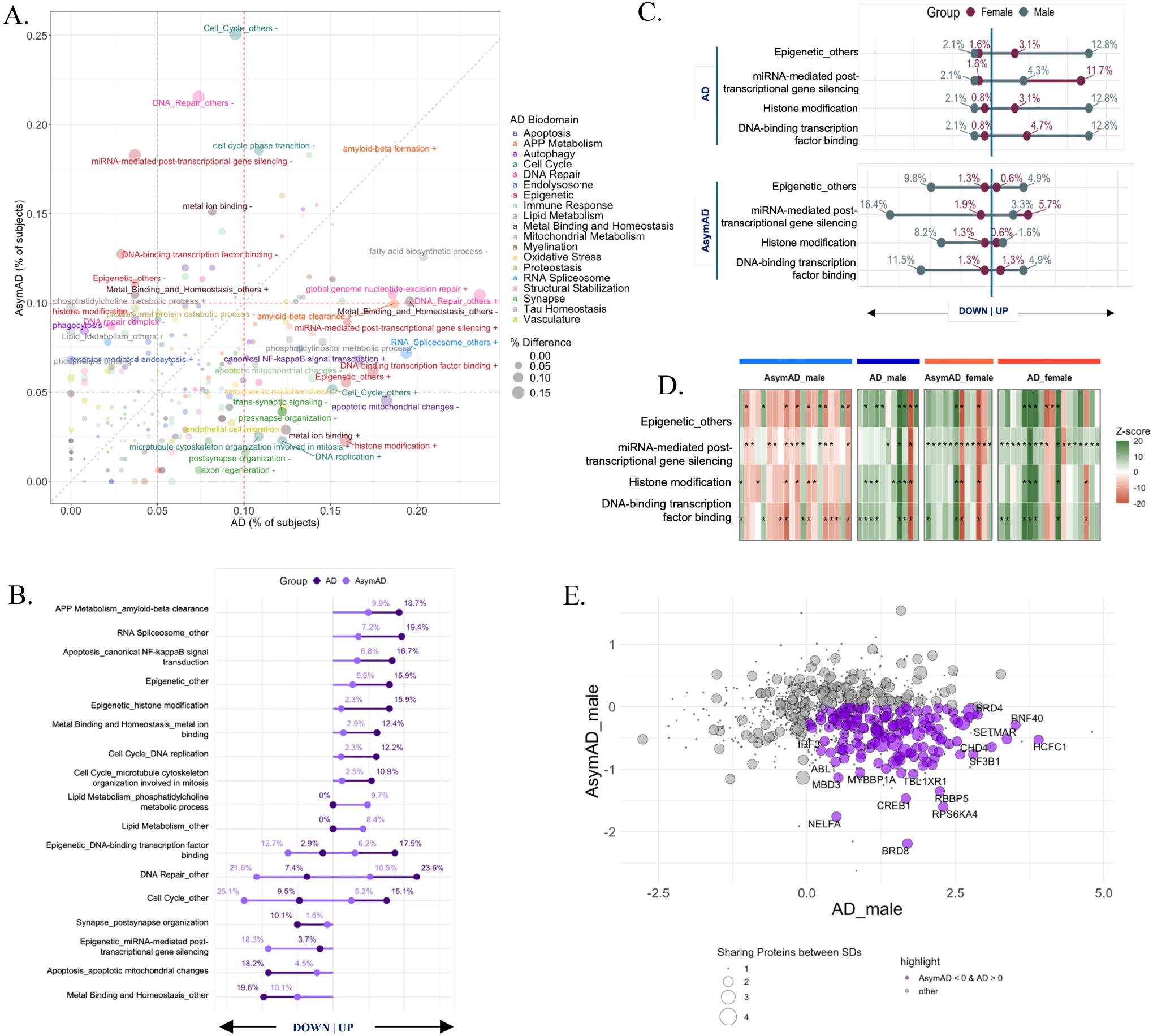
INDIGO reveals significant AD biodomain alterations with distinct patterns in AD and AsymAD groups. **(A)** Frequencies of significant biodomain and subdomain deviations in AsymAD and AD individuals, derived from INDIGO z-scores. The direction of dysregulation is indicated by “+” (up) or “–” (down) appended to subdomain labels. **(B)** Direction-separated frequencies of subdomains showing the largest differences between AD and AsymAD, highlighting multiple epigenetic subdomains with disease stage-dependent opposite dysregulation. **(C)** Sex-stratified enrichment frequencies of four epigenetic subdomains, showing that differences between AD and AsymAD are driven primarily by male subjects. **(D)** Individual-level z-scores for epigenetic subdomains. Significant alterations are marked with “*”. In males, dysregulation is distributed across individuals for transcription factor binding, histone modification, and Epigenetic_others subdomains with alterations of those subdomains are confined to few females. Consistent upregulation observed for miRNA-mediated post-transcriptional gene silencing in female subjects. **(E)** Average protein-level z-scores across male groups within epigenetic subdomains; proteins consistently upregulated in AD and downregulated in AsymAD across ≥2 subdomains are highlighted in purple. Proteins consistently upregulated in AD and downregulated in AsymAD males—and present in at least two epigenetic subdomains—were flagged as robust markers, including BRD8, CREB1, RBBP5, RPS6KA4/MSK2, CHD4, and SF3B1.

These epigenetic differences were primarily driven by males, with AD males exhibiting stronger activation than AsymAD males, whereas females showed more heterogeneous patterns (Figure 3C–D). Protein-level summaries supported coordinated shifts across epigenetic regulators during disease progression (Figure 3E).

Several subdomains also exhibited clear sex bias in AD, with female-enriched dysregulation involving amyloid-β formation, autophagy, and oxidative stress, and male-enriched dysregulation involving DNA repair, RNA splicing, and cell cycle–related processes (Figure S7). Notably, amyloid-β formation showed the strongest female-biased dysregulation, with female AD individuals exhibiting broader and more coordinated molecular alterations than males (Figure S8).

Together, these results identify epigenetic biodomains as prominent markers of the transition from asymptomatic to symptomatic AD and reveal sex-dependent functional organization that is consistent across pathway, subdomain, and biodomain levels.

### 3.4 Individual proteomic signatures correlate with clinical metrics

To link individualized molecular profiles with clinical markers of AD, we correlated INDIGO-derived subdomain z-scores with APOE4 allele dosage, Braak stage, CERAD score, and MMSE across the entire study cohort, including controls (Figure 4A). This approach captures continuous variation across the clinical spectrum rather than restricting analysis to categorical disease stages, thereby enhancing resolution of clinicopathological associations.

**Figure 4.**
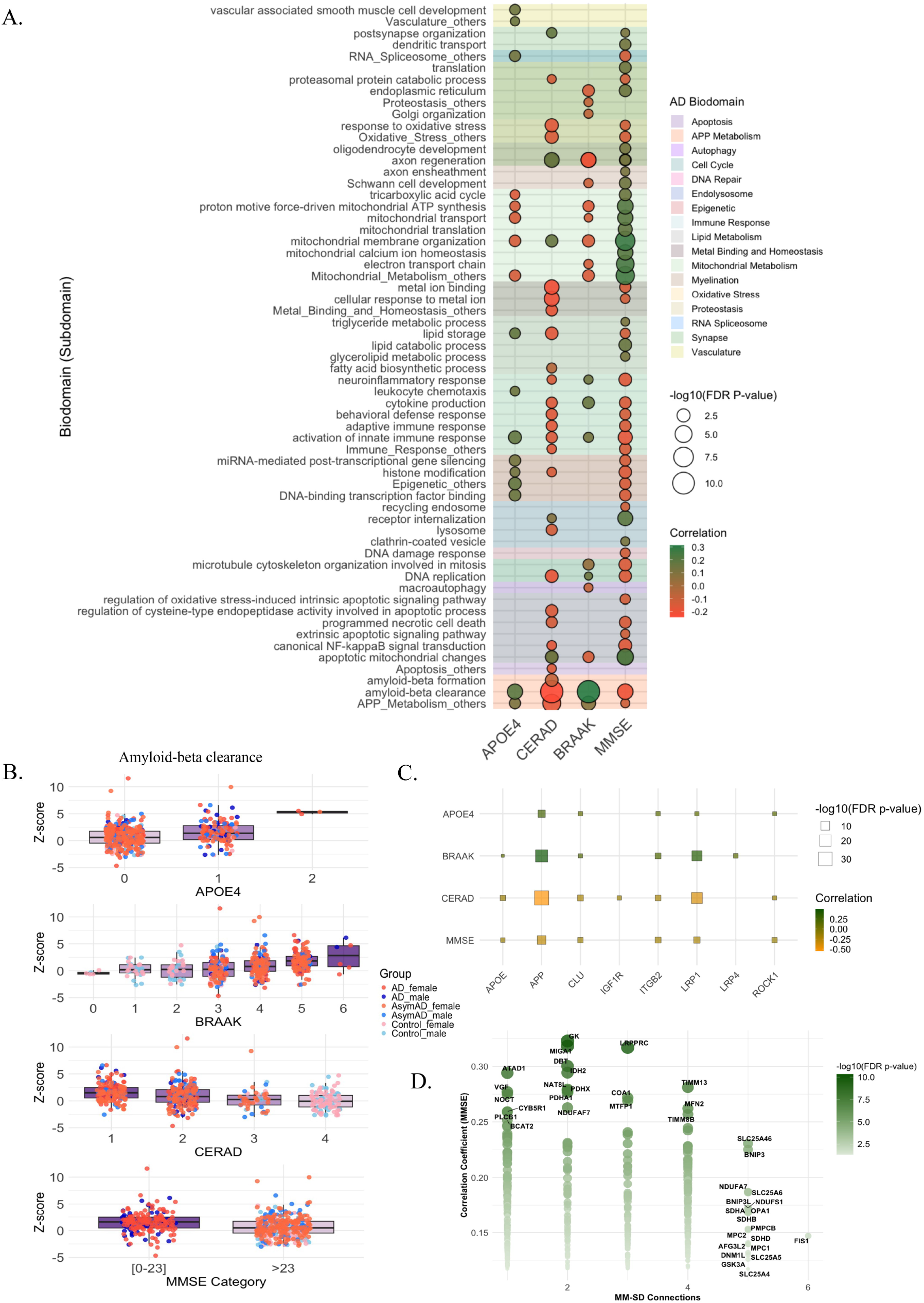
Subdomain-Level Proteomic Signatures Linked to Cognitive and Neuropathological Metrics. (A) Heatmap showing correlation between INDIGO-derived subdomain z-scores and four clinical and pathological variables—APOE4 dosage, Braak stage, CERAD score, and MMSE—computed across all subjects. Positive correlations with Braak stage or APOE4 indicate increased subdomain activity with greater disease burden or genetic risk, whereas positive correlations with CERAD or MMSE indicate higher subdomain activity associated with reduced amyloid pathology or better cognitive performance. **(B)** Stratified distributions of amyloid-β clearance subdomain scores, showing progressive increases with higher APOE4 dosage, greater pathology burden and worsening cognition. **(C)** Correlation plots for representative amyloid clearance proteins demonstrating links to amyloid, tau, and cognition. At the molecular level, 8 of 38 proteins within this subdomain were significantly correlated with clinical metrics. These proteins converge on pathways central to amyloid clearance, lipid trafficking, and synaptic integrity. Several (APOE, APP, CLU, PICALM) are established AD risk genes, LRP1/4 regulate receptor-mediated Aβ clearance, and ROCK1 promotes tau phosphorylation and synaptic degeneration. **(D)** Network-style visualization highlighting shared and subdomain-specific mitochondrial proteins associated with MMSE. In total, 2,196 non-overlapping proteins spanning twelve mitochondrial metabolism subdomains (MM-SDs) were evaluated for associations with four clinical variables— APOE4 dosage, Braak stage, CERAD score, and MMSE—across all subjects. Of the 495 proteins associated with MMSE, 374 exhibited significant positive correlations after multiple-testing correction (p-adjusted < 0.05), indicating that preserved or elevated mitochondrial protein activity is broadly linked to better cognitive performance. Proteins with the strongest correlation coefficients were often confined to fewer MM-SDs, suggesting specialized roles within discrete mitochondrial processes. In contrast, proteins exhibiting moderate but reproducible correlations were more frequently shared across multiple mitochondrial subdomains, highlighting a set of integrative regulators spanning mitochondrial transport, membrane organization, and metabolic coupling. Proteins shared across four to six MM-SDs, including FIS1 and members of the SLC family (SLC25A4, SLC25A5, SLC25A6), positioning them as hub-like components of a common mitochondrial axis associated with cognitive resilience.

Multiple subdomains exhibited coherent and clinically meaningful associations across all four clinical measures (Figure 4A). Subdomains related to amyloid-β clearance and innate immune activation showed consistent positive correlations with APOE4 dosage and Braak stage, alongside negative correlations with CERAD score and MMSE, indicating increased activation of these processes with greater amyloid and tau pathology and worsening cognition. In contrast, mitochondrial membrane organization, electron transport, and mitochondrial translation, displayed the opposite pattern, with negative correlations to Braak stage and positive correlations to MMSE, consistent with preserved mitochondrial function being associated with lower tau burden and better cognitive performance. Interestingly, dendritic transport, mitochondrial translation, and lipid catabolic processes, were positively correlated with MMSE independent of amyloid or tau pathology, suggesting distinct pathways may sustain cognitive resilience even in presence of neuropathological burden. Indeed, recent literature have shown that preservation of dendritic spine structure is strongly associated with cognitive resilience^25, 26^. Together, these correlation patterns define opposing functional axes aligned with disease burden versus cognitive preservation across the AD continuum.

The *APP metabolism—amyloid-*β *clearance* subdomain showed one of the strongest disease-linked signatures, with higher scores associated with increased APOE4 dosage, greater amyloid and tau pathology, and worse cognitive performance (Figure 4A–B). Protein-level analyses within this subdomain identified a subset of proteins whose expression tracked consistently with clinical measures, including several established AD risk genes and regulators of amyloid trafficking and synaptic integrity (Figure 4C). These results demonstrate that the amyloid clearance axis integrates genetic susceptibility, amyloid and tau, and cognitive decline, validating INDIGO’s capacity to resolve clinically meaningful proteomic signals.

In contrast, mitochondrial membrane organization defined a putative protective axis. Higher subdomain scores were associated with lower Braak stage and better cognitive performance (Figure 4A). A large fraction of mitochondrial proteins showed positive associations with MMSE, indicating a widespread link between preserved mitochondrial function and cognitive resilience (Figure 4D). Proteins shared across multiple mitochondrial subdomains, including FIS1 and members of the SLC25 family, emerged as integrative components of this resilience-associated axis.

These results demonstrate that individualized functional signatures capture both pathogenic and protective molecular processes that align with clinical and neuropathological variation across the AD spectrum.

### 3.5 Graph-based clustering identifies molecularly distinct subgroups of AD and AsymAD subjects

To identify individuals with shared functional dysregulation patterns, we constructed a bipartite graph linking subjects to significantly altered subdomains and applied graph clustering to identify recurrent modules (Figure 5A). This analysis identified 29 clusters, of which nine were sufficiently prevalent to warrant detailed analysis (Figure 5B). These clusters revealed coordinated molecular programs that transcended diagnostic and sex categories (Figures 5C and S9). Among the nine prevalent clusters, several exhibited sharply contrasting patterns of subdomain enrichment, revealing molecularly defined subgroups not apparent from clinical classification alone.

**Figure 5.**
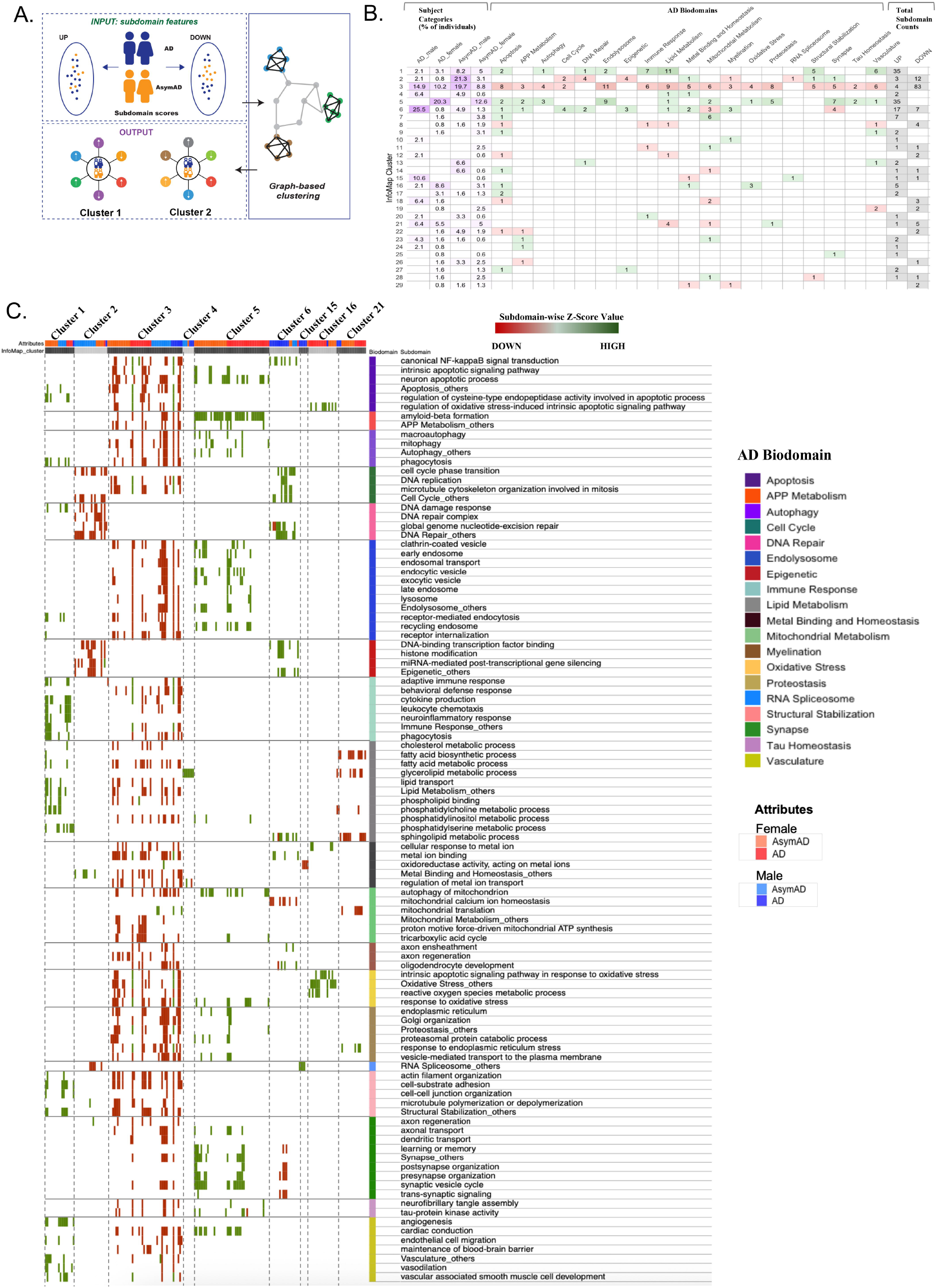
Graph-based clustering reveals molecularly distinct subgroups of AD and AsymAD. **(A)** Schematic of the bipartite network structure. Subdomains are shown as color-coded squares, with yellow and pink indicating up- and down-regulation, respectively. Subjects are represented by colored circles: blue (AD males), red (AD females), cyan (AsymAD males), and orange (AsymAD females). **(B)** The 29 clusters identified by the InfoMap algorithm and the distribution of subdomains and individual characteristics within the clusters. Nine clusters were sufficiently represented (≥10% of aggregated individuals in each cluster) for downstream analysis. **(C)** Network plots of the nine clusters that were sufficiently represented (≥5% of individuals) for downstream analysis. (D) Individual-level signatures defining major clusters, highlighting coordinated alterations in neuronal, mitochondrial, immune, vascular, and epigenetic processes.

Clusters 1 and 3 exhibited the strongest reciprocal contrast across four biodomains: immune response, lipid metabolism, structural stabilization, and vasculature (Figure 5C). Cluster 1 showed consistent upregulation across these modules, whereas Cluster 3 displayed coordinated downregulation. Notably, Cluster 3 also demonstrated broader suppression of synaptic, mitochondrial, and proteostasis-related subdomains and included both AD and AsymAD individuals, suggesting that this molecular configuration may emerge prior to clinical conversion and persist across disease stages.

A second reciprocal pattern was observed between Clusters 2 and 6, driven by subdomains associated with cell cycle regulation, DNA repair, and epigenetic processes (Figures 5C and S10A). These modules were uniformly downregulated in Cluster 2, which was enriched for AsymAD individuals, and upregulated in Cluster 6, which was enriched for AD subjects. Although both clusters were male-dominated, they diverged sharply in regulatory direction, highlighting stage-dependent alterations within genome maintenance pathways.

Two clusters showed marked sex specificity. Cluster 5 consisted exclusively of female individuals and was characterized by coordinated upregulation of amyloid-β formation, autophagy, endolysosomal trafficking, proteostasis, and synaptic subdomains. Cluster 16, another female-dominated subgroup, displayed a focused and consistent upregulation of oxidative stress–related subdomains, defining a distinct molecular trajectory within a subset of females.

Protein-level analyses supported these cluster-level distinctions. Differential expression between Clusters 1 and 3 identified coherent sets of proteins spanning the four discriminating biodomains (Figure 6A–B and Figure S11A), while comparison of Clusters 2 and 6 revealed divergent regulation of proteins involved in epigenetic and cell-cycle control (Figure S10A-E). In Cluster 16, oxidative stress–related proteins showed consistent elevation, with PARK7 exhibiting the strongest and most specific increase (Figure 6C–E and S11B-C).

**Figure 6.**
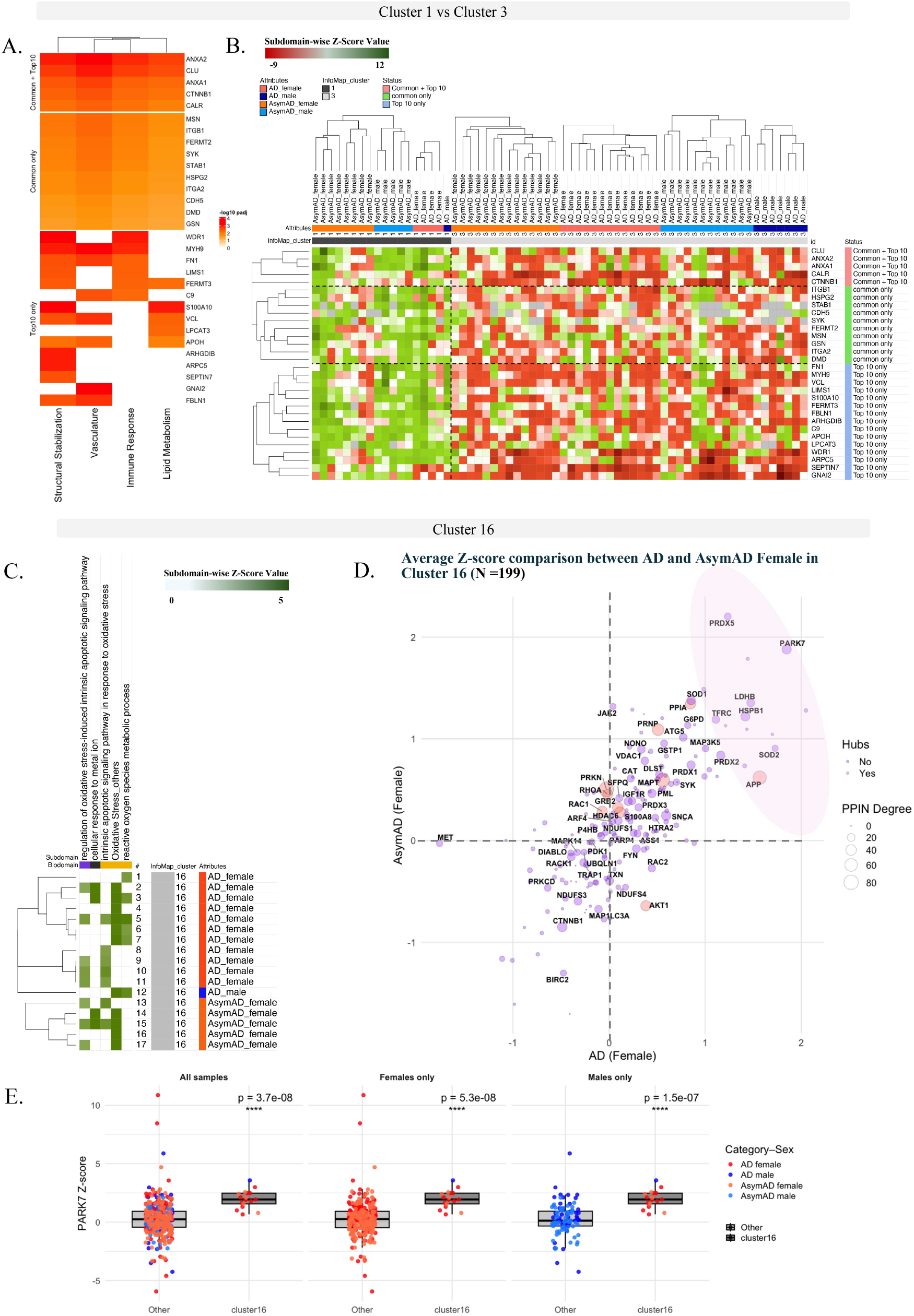
Protein-level characterization of key clusters from graph-based analysis. (A) Heatmap summarizing the results of protein-wise Wilcoxon rank-sum tests comparing Cluster 1 and Cluster 3 across the four biodomains that most strongly distinguish the two groups. Proteins are highlighted: (i) those common to all four biodomains (“Common only”), (ii) the top 10 most significantly different proteins within each biodomain (“Top10 only”), and (iii) proteins appearing in both categories (“Common + Top10”). Rows correspond to protein name and columns represent the four biodomains (structural stabilization, lipid metabolism, immune response, and vasculature). Cell color reflects the –log10(P value) of the statistical difference between clusters. (B) Heatmap of z-score profiles for proteins selected from panel A, including (i) proteins common across all four biodomains and (ii) the top 10 most significantly different proteins within each biodomain. Values are shown for all individuals in Clusters 1 and 3, illustrating the coordinated patterns of up- and downregulation across clusters. (C) Subdomain-specific z-score profiles for Cluster 16 individuals across five curated subdomains, including three oxidative stress–related modules. (D) Scatter plot of average protein z-scores (199 oxidative-stress and related proteins) comparing AD vs. AsymAD females within Cluster 16. Proteins exhibiting concordant upregulation in both groups are highlighted. Several proteins showed concordant elevation in both AD and AsymAD females, with PARK7 demonstrating the most robust increase. (E) Boxplots comparing PARK7 z-scores in Cluster 16 versus all other samples, stratified by sex and diagnosis. Elevated PARK7 levels are highly specific to Cluster 16 females.

Finally, we evaluated whether cluster membership aligned with clinical or neuropathological measures. Clinical association analyses indicated that Cluster 2 was positively associated with MMSE, whereas Cluster 16 was associated with advanced age at death (Figure S12A-B). Collectively, these results demonstrate that graph-based clustering of individualized functional profiles resolves biologically coherent molecular subgroups across the AD continuum.

## 4 DISCUSSION

A major challenge in AD research is the extensive inter-individual molecular heterogeneity that is not captured by traditional case–control DE analyses. Precision medicine efforts in AD aim to move beyond population-averaged, symptom-based approaches toward biomarker-guided strategies tailored to disease-stage and individual biology^27^. While N-of-1 methodologies have been increasingly applied in prognostic modeling^28^, clinical trials^29, 30^, and genetic risk assessment^31^, their translation to therapeutic discovery in AD remains limited. This gap reflects both the complexity of aging-related neurodegeneration and the difficulty of extracting robust individual-level signals from high-dimensional omics data. In this study, we present INDIGO, a proof-of-concept framework that adapts precision omics strategies to enable systematic quantification of single-subject proteomic deviations across AD continuum and their integration into biologically informed functional contexts.

Aggregating individualized proteomic deviations into pathways and AD-specific biodomains revealed sex-stratified and stage-dependent patterns of dysregulation that are hidden in cohort-level analyses. Consistent with growing evidence of sex differences in AD^32, 33^, we observed male-biased activation of epigenetic, DNA repair, and RNA-processing modules, contrasted with female-enriched dysregulation of mitochondrial, autophagic, and amyloid-associated pathways. These findings reiterate that males and females may engage distinct molecular programs in response to AD pathology, with implications for sex-informed therapeutic strategies.

A particularly striking result was the opposing regulation of epigenetic, cell-cycle, and DNA-repair biodomains between AsymAD and AD. These modules were predominantly downregulated in AsymAD but strongly upregulated in AD, suggesting a stage-dependent inversion of genome-maintenance and regulatory programs. Such patterns are consistent with prior reports linking chromatin remodeling, transcriptional dysregulation, and aberrant cell-cycle re-entry to AD progression and neuronal vulnerability^34-36^. Importantly, this pattern was independently recapitulated by graph-based clustering, where a predominantly AsymAD male cluster (Cluster 2) exhibited coordinated downregulation of these pathways, while an AD-enriched male cluster (Cluster 6) showed upregulation. Together, these multilevel observations suggest a stage-associated shift from relatively compensated or quiescent regulatory states in AsymAD to dysregulated states in overt AD.

A key strength of INDIGO is its ability to link functional signatures to continuous clinical and neuropathological measures, enabling biologically interpretable connections between molecular profiles and disease severity. Multiple subdomains—including innate immune activation, amyloid processing, lipid metabolism, and mitochondrial organization—showed robust associations with Braak stage, CERAD score, MMSE, and APOE4 dosage. Notably, the amyloid-β clearance subdomain exhibited coherent associations with APOE4 dosage, tau and amyloid pathology, and cognitive impairment, reinforcing its centrality in AD biology^37, 38^. In contrast, mitochondrial membrane organization emerged as a putative protective axis, consistent with recent evidence linking preserved mitochondrial function to cognitive resilience^39^. These results highlight how individualized functional profiling can inform both pathogenic and resilience-related mechanisms across preclinical and symptomatic stages.

Finally, graph-based clustering identified molecularly coherent subgroups that transcend conventional diagnostic categories and captured structured biological heterogeneity within AD. Clusters defined by immune–vascular activation, synaptic–mitochondrial suppression, genome-maintenance dysregulation, or oxidative stress showed distinct clinical associations, including differences in cognition and age at death. Such molecular stratification extends emerging multi-omic subtyping efforts and suggests a robust patient classification for clinical trials and targeted interventions. Additional subtyping frameworks based on neuropathology^40^, cerebrospinal fluid (CSF) proteomics^41, 42^, and cognitive profiles^43^ have also revealed distinct AD trajectories. INDIGO offers a potential bridge between these modalities by integrating individual-level molecular states with multi-domain clinical subtypes.

While the findings provide new insights in AD heterogeneity, some limitations warrant consideration. Our analyses rely on cross-sectional postmortem proteomic data, limiting temporal inference and direct assessment of individual disease trajectories. Sample sizes in some sex- and diagnosis-stratified groups were modest, and functional interpretations depend on existing pathway and biodomain annotations. Future studies incorporating longitudinal designs, multi-regional and multi-omic data, and independent validation cohorts will be essential. Nevertheless, by integrating single-subject deviation profiling, functional aggregation, clinical correlation, and graph-based stratification, we provide a scalable systems-level framework for advancing precision medicine strategies in AD.

## Acknowledgements/AI Disclosure

The results published here are in whole or in part based on data obtained from the AD Knowledge Portal (https://adknowledgeportal.org). Data used in this study were obtained from the Accelerating Medicines Partnership Program for Alzheimer’s Disease (AMP□AD) Consortium members below: Mayo RNAseq Study: Study data were provided by the following sources: The Mayo Clinic Alzheimer’s Disease Genetic Studies, led by Dr. Nilufer Ertekin□Taner and Dr. Steven G. Younkin, Mayo Clinic, Jacksonville, FL, using samples from the Mayo Clinic Study of Aging, the Mayo Clinic Alzheimer’s Disease Research Center, and the Mayo Clinic Brain Bank. Data collection was supported through funding by NIA grants P50 AG016574, R01 AG032990, U01 AG046139, R01 AG018023, U01 AG006576, U01 AG006786, R01 AG025711, R01 AG017216, R01 AG003949, NINDS grant R01 NS080820, CurePSP Foundation, and support from Mayo Foundation. Study data include samples collected through the Sun Health Research Institute Brain and Body Donation Program of Sun City, Arizona. The Brain and Body Donation Program is supported by the National Institute of Neurological Disorders and Stroke (U24 NS072026 National Brain and Tissue Resource for Parkinson’s Disease and Related Disorders), the NIA (P30 AG19610 Arizona Alzheimer’s Disease Core Center), the Arizona Department of Health Services (contract 211002, Arizona Alzheimer’s Research Center), the Arizona Biomedical Research Commission (contracts 4001, 0011, 05□901, and 1001 to the Arizona Parkinson’s Disease Consortium), and the Michael J. Fox Foundation for Parkinson’s Research. Religious Orders Study/Memory and Aging Project (ROSMAP): We are grateful to the participants in the Religious Order Study and the Memory and Aging Project. This work was supported by the US National Institutes of Health (U01 AG046152, R01 AG043617, R01 AG042210, R01 AG036042, R01 AG036836, R01 AG032990, R01 AG18023, RC2 AG036547, P50 AG016574, U01 ES017155, KL2 RR024151, K25 AG041906□01, R01 AG30146, P30 AG10161, R01 AG17917, R01 AG15819, K08 AG034290, P30 AG10161, and R01 AG11101). Mount Sinai Brain Bank (MSBB): This work was supported by grants R01AG046170, RF1AG054014, RF1AG057440, and R01AG057907 from the NIH/NIA. R01AG046170 is a component of the AMP□AD Target Discovery and Preclinical Validation Project. Brain tissue collection and characterization was supported by NIH HHSN271201300031C.

The authors acknowledge the use of generative artificial intelligence tools to assist with language editing, manuscript organization, and refinement of scientific phrasing. All scientific content, data analyses, interpretations, and conclusions were conceived, validated, and approved by the authors, who take full responsibility for the accuracy and integrity of the work. No generative AI tools were used to generate data, perform analyses, or make scientific decisions.

## Funding Sources

This work was supported by the National Institute of Health grants U54 AG054345 and U54 AG065187.

## Conflict of Interest

The authors declare that this research was conducted in the absence of any commercial or financial relationships that could be construed as a potential conflict of interest.

## Consent Statement

No consent was required as all human subjects data were reused under controlled access with an active AD Knowledge Portal Data Use Certificate (v7.3). All data were anonymized by original sources with no possibility of deanonymization.

## Notes

https://avijitpodder-ap.github.io/ssproteomics-workflow/

### Competing Interest Statement

The authors have declared no competing interest.

