## Supplemental materials for "Single-subject Proteomic Signatures in Alzheimer’s Disease Reflect Clinical Phenotypes and Distinguish Asymptomatic from Symptomatic Cases"

- **Supplementary Methods**

### 2.1 Proteomic preprocessing and quality control

Proteomic data underwent normalization using pooled-reference scaling and batch-effect correction with TAMPOR, as provided by the ROSMAP proteomics pipeline (ROSMAP Proteomics Data and Methods- https://help.adknowledgeportal.org/apd/Proteomics-Data-and-Methods.2826010634.html). Proteins with >50% missing values were excluded prior to downstream analysis. No retained sample exhibited >20% missingness across proteins (Figure S1A).

To address redundancy from repeated protein identifiers, protein entries were collapsed by gene symbol, retaining the instance with the highest absolute row-sum across all individuals as the representative profile (Figure S1B). After preprocessing, each individual had quantified abundances for approximately 7,000–8,000 proteins.

Additional details on data processing and quality control are available via the AD Knowledge Portal documentation.

### 2.2 ssDEA reference construction and quality control

To establish a stable and unbiased reference for ssDEA, we explicitly evaluated within-group variability among control samples. Given that z-score–based deviation analysis is sensitive to heterogeneity in the reference population, uncontrolled variability or outlier control profiles could introduce bias or inflate apparent deviations in disease subjects. We therefore implemented a leave-one-out (LOO) z-score procedure, in which each control subject was iteratively excluded and treated as a test sample, and protein-wise z-scores were computed relative to the remaining sex-matched controls. The resulting LOO z-score matrices were subjected to hierarchical clustering to identify controls exhibiting systematic deviations from the expected reference structure. Control samples that consistently segregated as outliers or formed distinct branches in the dendrogram were removed prior to final reference construction (Figure S2A–D). This procedure yielded a refined, internally consistent control cohort, thereby improving the accuracy and interpretability of individual-level deviation estimates in AD and AsymAD subjects (Figure S3A–D).

**2.3 Cohort-based differential expression comparison**

Cohort-based differential expression results were derived from previously published ROSMAP TMT proteomics analyses comparing AD cases with controls (Synapse ID: syn25607662). Protein rankings were based on multiple-testing–adjusted p-values. Individual ssDEA rankings were derived from absolute z-score magnitude. Overlap was computed per subject and summarized across groups.

**2.4 ssFPA score computation**

For each subject $j$, overlap between the quantified proteome and each functional module was determined, and protein-level z-scores were aggregated using Stouffer's Z method.

For a pathway or subdomain $P$ containing k proteins with non-missing z-scores, the functional score was calculated as:

$$Z_P^{(j)} = \frac{1}{\sqrt{k}} \sum_{i=1}^{k} z_i^{(j)} \qquad \text{where } z_i \neq \text{NA}$$

where $z_i^{(j)}$ denotes the z-score for protein $i$ in subject $j$. Missing z-scores were excluded prior to aggregation to avoid bias from imputation and ensure robustness of subject-level functional scores.

KEGG pathway annotations [1] were retrieved using the KEGGREST R package (v1.48.1). The AD-specific biodomain [2] and subdomain annotations were obtained from Synapse (ID: syn26529354).

#### 2.5 Reference-based functional significance thresholds

Functional perturbations were defined relative to control-derived empirical distributions rather than fixed thresholds to account for pathway-specific variability and proteomic noise. The 1st–99th percentile bounds capture expected non-pathological variation and enable sensitive detection of subtle yet biologically meaningful deviations across disease stages.

#### 2.6 Statistical framework for clinical associations

AD and AsymAD subjects were analyzed jointly to reflect the continuum model of AD progression, in which AsymAD represents an intermediate or prodromal state. For ordinal variables (APOE4 dosage, Braak stage, CERAD score), Kruskal–Wallis tests were used, complemented by linear models to assess trend effects. Pearson correlation was used to evaluate associations between MMSE and subdomain-level z-scores. Subdomains with adjusted p-values < 0.05 for any metric were retained for downstream interpretation.

#### 2.7 Graph construction and community detection

An undirected bipartite graph *G=(V, E)* was constructed, where vertices included all AD and AsymAD subjects and all functional subdomains. An edge was drawn between subject $j$ and subdomain $s$ if the subject exhibited a z-score outside the control-derived 1st–99th percentile range for that subdomain. z-scores were transformed to absolute values to reflect the magnitude of dysregulation irrespective of direction, and all edges were treated as unweighted.

Communities were identified using the Infomap algorithm, which detects modules by minimizing the description length of random walks on the graph [3]. The objective function minimized by Infomap is given by:

$$L = \ q_{\curvearrowright} H(Q) + \sum_{i=1}^{m} p_i H(P_i)$$

where $q_{\curvearrowright}$ denotes the probability of exiting any module, $\mathrm{H}(Q)$ is the entropy of movements between modules, $p_i$ is the probability of remaining within module $i$, $\mathrm{H}(P_i)$ is the entropy within module $i$, and $m$ is the total number of modules. Infomap's flow-based formulation is well suited for identifying fine-grained communities in bipartite graphs with heterogeneous connectivity.

### 2.8 Statistical testing of cluster-associated clinical features

Clinical and pathological variables were selected to capture genetic risk, disease severity, cognitive status, and technical covariates. Wilcoxon rank-sum (Mann–Whitney U) tests were used for pairwise comparisons between clusters, and adjusted p-values < 0.05 were considered statistically significant.

### 2.9 PPI network construction and protein prioritization

Protein–protein interaction (PPI) data were obtained from the Integrated Interactions Database (IID; version 2025-25), which contains over 1.5 million experimentally validated interactions across tissues and diseases [4]. Interactions were filtered to retain brain-specific, experimentally supported physical interactions, excluding predicted or orthology-based edges.

A brain-specific PPI network was constructed with nodes representing subdomain-associated proteins and edges representing high-confidence interactions. Network topology was analyzed to identify hub proteins based on degree centrality.

To integrate expression and network information, average protein-level z-scores were computed within diagnostic or cluster-defined groups. For each protein $p$ in group $g$, the average z-score was calculated as:

$$\bar{Z}_p^{(g)} = \frac{1}{n_g} \sum_{i=1}^{n_g} Z_{p,i}$$

where $Z_{p,i}$ denotes the z-score of protein p in subject i, and $n_g$ is the number of individuals with non-missing values for that protein. Highly connected proteins exhibiting consistent up- or downregulation were prioritized as candidate regulators of subdomain-level dysregulation.

#### 2.10 Software and visualization tools

Analyses were conducted in R (v4.5.0). ROSMAP data were accessed using the synapser package [5]. KEGG pathway annotations were retrieved using KEGGREST, and AD subdomain mappings were implemented using in-house functions [6]. Graph construction and community detection were performed with igraph and the Infomap algorithm [7]. Data visualization utilized ggplot2 and patchwork for statistical graphics [8, 9], Morpheus for heatmaps (https://software.broadinstitute.org/morpheus), and Cytoscape for protein–protein interaction network visualization [10].

### Supplementary Methods References

[1] Kanehisa M, Goto S. KEGG: kyoto encyclopedia of genes and genomes. Nucleic Acids Res. 2000;28:27-30.

[2] Cary GA, Wiley JC, Gockley J, Keegan S, Amirtha Ganesh SS, Heath L, et al. Genetic and multi-omic risk assessment of Alzheimer's disease implicates core associated biological domains. Alzheimers Dement (N Y). 2024;10:e12461.

[3] Rosvall M, Bergstrom CT. Maps of random walks on complex networks reveal community structure. Proc Natl Acad Sci U S A. 2008;105:1118-23.

[4] Kotlyar M, Pastrello C, Sheahan N, Jurisica I. Integrated interactions database: tissue-specific view of the human and model organism interactomes. Nucleic Acids Res. 2016;44:D536-41.

[5] Hoff B. synapser: R language bindings for Synapse API. 0.1.7 ed: Sage-Bionetworks; 2017.

[6] Tenenbaum D, Maintainer B. KEGGREST: Client-side REST access to the Kyoto Encyclopedia of Genes and Genomes (KEGG). R package version 1.48.1; 2025.

[7] Csárdi GN, T.; Traag, V.; Horvát, S.; Zanini, F.; Noom, D.; Müller, K. igraph: Network Analysis and Visualization in R. 2.1.4 ed: The R Foundation for Statistical Computing; 2025.

[8] Wickham H. ggplot2: Elegant Graphics for Data Analysis: Springer-Verlag; 2016.

[9] Pedersen TL. patchwork: The Composer of Plots. 1.2.0 ed: The R Foundation for Statistical Computing; 2024.

[10] Shannon P, Markiel A, Ozier O, Baliga NS, Wang JT, Ramage D, et al. Cytoscape: a software environment for integrated models of biomolecular interaction networks. Genome Res. 2003;13:2498-504.

- **Supplemental Figure Legends**

**Suppl. Fig. 1. Quality control assessment of missing values and handling of repeated protein identifiers.**

(A) Distribution of missing protein abundance values per individual across all diagnostic and sex groups. Each bar represents a single subject, ordered by category (Control, AD, AsymAD, and excluded individuals). The plot illustrates substantial heterogeneity in missingness across samples, with some individuals exhibiting >1,500 missing protein measurements. To ensure robustness in downstream analyses, we applied a missingness filter and retained only individuals with ≤20% missing proteins across the full panel.

(B) Frequency of repeated Protein_name identifiers prior to preprocessing. Several proteins appeared multiple times in the dataset (e.g., HLA-A with 5 entries, HLA-B with 4 entries, and others with 2 entries each), reflecting redundancy arising from different peptide or isoform measurements. To resolve this duplication, repeated entries were collapsed by selecting the instance with the highest absolute row-sum across all individuals, yielding a single representative abundance profile for each protein. This ensured that all proteins included in subsequent analyses corresponded to unique, non-redundant identifiers.

**Suppl. Fig. 2. Leave-One-Out stability assessment of protein-level z-score distributions in control subjects.**

Heatmaps showing the distribution of protein-wise z-scores computed using a Leave-One-Out (LOO) procedure across all control females (A) and control males (B). Hierarchical clustering of LOO-normalized z-scores revealed one outlier sample in each group (far left in both heatmaps), characterized by disproportionate branching and separation in the dendrogram. These dendrogram outlier indicated potential sample-specific deviation within the control background distribution.

LOO analysis repeated after removing the identified outgrowth samples for control females (C) and control males (D). The resulting heatmaps show stable, homogeneous clustering across individuals, with no further irregular branching patterns. z-score distributions remain tightly centered around zero, indicating that the remaining controls form a consistent and reliable reference set.

**Suppl. Fig. 3. Protein z-score–based clustering of AD and AsymAD subjects.**

Heatmaps showing hierarchical clustering of AD females (A), AD males (B), AsymAD females (C), and AsymAD males (D) based on protein-level z-scores. Rows represent proteins and columns represent individual subjects within each diagnostic/sex subgroup; dendrograms are shown for both dimensions. The predominantly yellow color distribution indicates that z-scores remain centered around zero across individuals, with scattered regions of deviations (green = positive Z, red = negative Z) that vary between individuals.

**Suppl. Fig. 4. Expanded assessment of ssDEA-derived protein rankings beyond the top 50 proteins in AD males.**

(A) Heatmap showing the ssDEA-based ranking of significant cohort-level proteins (Cor_Pval < 0.05) across AD male individuals. Each cell encodes the rank of a protein within a subject, with red indicating proteins appearing in the top 50 for that subject. Although many proteins are consistently significant at the cohort level, only a subset appears frequently among the highest-ranked proteins in single-subject analyses, illustrating substantial inter-individual variability.

(B) Frequency matrix summarizing how often each protein appears within expanded rank thresholds—top 10, top 50, top 100, and top 500—across AD male subjects. This analysis extends the top-50 evaluation and demonstrates that increasing the rank window does not alter the overall pattern: only a minority of cohort-significant proteins occur repeatedly among high-ranking positions across individuals, whereas most proteins show limited recurrence even at thresholds as broad as top 500.

(C) Spearman correlation matrix evaluating concordance among rank-frequency groups (top 10, 50, 100, 500). Each cell represents the correlation between two ranking thresholds (e.g., top 10 vs. top 50). Strong positive correlations, particularly among adjacent thresholds, indicate consistent relative ranking behavior across subjects. This further supports that individual-level protein rankings maintain substantial granularity even when moving beyond the top 50.

**Suppl. Fig. 5. Expanded assessment of ssDEA-derived protein rankings beyond the top 50 proteins in AD females.**

(A) Heatmap showing the ssDEA-based ranking of significant cohort-level proteins (Cor_Pval < 0.05) across AD female individuals. Each cell encodes the rank of a protein within a subject, with red indicating proteins appearing in the top 50 for that subject. Although many proteins are consistently significant at the cohort level, only a subset appears frequently among the highest-ranked proteins in single-subject analyses, illustrating substantial inter-individual variability.

(B) Frequency matrix summarizing how often each protein appears within expanded rank thresholds—top 10, top 50, top 100, and top 500—across AD female subjects. This analysis extends the top-50 evaluation and demonstrates that increasing the rank window does not alter the overall pattern: only a minority of cohort-significant proteins occur repeatedly among high-ranking positions across individuals, whereas most proteins show limited recurrence even at thresholds as broad as top 500.

(C) Spearman correlation matrix evaluating concordance among rank-frequency groups (top 10, 50, 100, 500). Each cell represents the correlation between two ranking thresholds (e.g., top 10 vs. top 50). Strong positive correlations, particularly among adjacent thresholds, indicate consistent relative ranking behavior across subjects. This further supports that individual-level protein rankings maintain substantial granularity even when moving beyond the top 50.

**Suppl. Fig. 6. Sex- and diagnosis-stratified frequencies of upregulated and downregulated AD subdomains.**

(A) Heatmap showing the proportion of individuals within each group (AD females, AsymAD females, AD males, AsymAD males) exhibiting upregulation of AD-related subdomains. Subdomains (rows) are ordered by hierarchical clustering based on their upregulation profiles. Values represent the fraction of subjects in each group with a positive subdomain z-score.

(B) Heatmap showing the proportion of individuals in each group exhibiting downregulation of AD subdomains. Rows are hierarchically clustered to illustrate shared and distinct suppression patterns across sex and diagnosis categories.

**Suppl. Fig. 7. Sex-specific patterns of subdomain dysregulation within AD subjects.**

(A) Scatter plot showing the percentage of AD males (x-axis) and AD females (y-axis) exhibiting deregulation for each AD subdomain. Each point represents a subdomain, colored by biodomain membership. The position in the plot indicates sex bias in deregulation frequency: points above the diagonal reflect female-enriched dysregulation, whereas points to the right of the vertical reference line indicate male-enriched dysregulation. Circle size corresponds to the absolute difference in deregulation frequency between sexes.

(B) Bar plot summarizing the direction and magnitude of sex differences for each dysregulated subdomain in AD subjects. Each row represents an AD subdomain, upregulated (right) and downregulated (left). The bar lengths indicate the percentage of male or female AD subjects (distinguished by color) exhibiting deregulation of that subdomain in the specified direction.

**Suppl. Fig. 8. amyloid-β–related molecular signatures across AD, AsymAD, and control subjects.**

(A) and (B) Density plot showing the distribution of subdomain-specific z-scores for female (A) and male (B) subjects across control (CT), AsymAD, and AD groups. Vertical dashed lines indicate significant single-subject deviations (beyond the control-derived 1st–99th percentile range) in AD and AsymAD individuals, stratified by sex. Rug plots below the density curves represent subdomain-level z-scores for individual subjects in each diagnostic category.

(C) Venn diagram highlighting the overlapping proteins present in the TMT proteomics assay from the amyloid-β formation subdomain.

(D) Heatmap of gene-wise z-scores for amyloid-β–related proteins across all individuals. Rows correspond to genes and columns to subjects, grouped by diagnosis and sex. Color intensity reflects the direction and magnitude of deviation (green, upregulation; brown, downregulation). Red asterisks denote proteins consistently deregulated across a large fraction of AD subjects, emphasizing coordinated pathway-level disruption.

(E) Functional protein–protein interaction (PPI) network constructed from amyloid-β–associated proteins. Nodes represent proteins and edges denote known functional interactions. Node size reflects network connectivity, highlighting APP as a central hub interacting with key AD-related proteins including APOE, CLU, BACE1, PSEN1/2, PRNP, and LRP family members, underscoring the coordinated dysregulation of amyloid-processing machinery in AD.

**Suppl. Fig. 9 – Selected graph-based clusters highlighting subject–subdomain relationships across AD and AsymAD.**

Network plots of the nine clusters that were sufficiently represented (≥10% of individuals) for downstream analysis. Nine representative InfoMap clusters (≥10% of aggregated individuals in each cluster) derived from the subject–subdomain bipartite network, illustrating coordinated patterns of functional dysregulation across individuals. Each panel corresponds to one cluster (Clusters 1, 2, 3, 4, 5, 6, 15, 16, and 21), depicting both subjects and enriched AD subdomains as nodes in a shared embedding space. Edges link individuals to the subdomains in which they exhibit significant up- or down-regulation. Node shape distinguishes AD and AsymaAD subjects from AD subdomains. Subject node coloring encodes diagnostic and sex groups (AD female, AD male, AsymAD female, AsymAD male) and AD subdomain node color indicates the direction of subdomain enrichment (UP or DOWN).

**Suppl. Fig. 10. Protein-level characterization of subdomain differences between Cluster 2 and Cluster 6.**

(A) Subdomain-wise z-score summary heatmap for Cluster 2 (AsymAD-enriched, predominantly male) and Cluster 6 (AD-enriched, predominantly male). Rows represent the seven subdomains associated with cell cycle regulation, DNA repair, and epigenetic processes; columns represent individuals from each cluster. Colors indicate direction of dysregulation (green = up, red = down). Cluster 2 shows consistent downregulation across these subdomains, whereas Cluster 6 shows widespread upregulation, aligning with the overall cluster-level contrast.

(B) Functional protein–protein interaction (PPI) network constructed for 1,435 curated non-overlapping proteins shared across the seven subdomains (1,420 proteins present in the PPIN). Nodes represent proteins, and node size reflects network degree (central connectivity).

(C) Scatter plot comparing average protein-level z-scores for 1,435 curated non-overlapping proteins across the seven subdomains. The x-axis shows the mean z-score in Cluster 6 (AD males), and the y-axis shows the mean z-score in Cluster 2 (AsymAD males). Proteins located in the lower-right quadrant (highlighted region) exhibit strong downregulation in Cluster 2 and concurrent upregulation in Cluster 6, representing key drivers of the contrasting subdomain patterns. Several proteins, including BRD4, EGFR, SNW1, and TRIM28, showed the strongest divergence between the two predominantly male clusters. Proteins annotated as PPIN hubs (top 1% based on degree, betweenness, and closeness centrality) are highlighted in red, emphasizing highly connected regulatory proteins contributing to the molecular divergence between clusters.

(D) Boxplots showing z-score distributions for selected proteins with strong opposing regulation between Cluster 2 and Cluster 6. Each subplot corresponds to a single protein, with individuals colored by diagnosis and sex. These proteins were chosen based on their effect sizes and network relevance and illustrate consistent downregulation in Cluster 2 and upregulation in Cluster 6.

(E) Heatmap summarizing the mapping of key differentiating proteins (from panel D) back to AD subdomains. Rows represent proteins, and columns represent subdomains within the three biodomains analyzed. Purple shading indicates presence of a protein within a given subdomain, demonstrating that the highlighted protein set spans multiple regulatory modules implicated in the Cluster 2 vs. Cluster 6 contrast.

**Suppl. Fig. 11. Identification and characterization of key proteins in the clusters.**

(A) Summary of protein-wise Wilcoxon rank-sum tests comparing Cluster 1 vs. Cluster 3 across the four biodomains that most strongly distinguished the two groups (immune response, lipid metabolism, structure stabilization, and vasculature). 486 proteins with adjusted $P < 0.05$ are highlighted.

(B) Functional protein–protein interaction (PPI) network constructed for 183 (out of 199) proteins associated with oxidative stress, mitochondrial function, proteostasis, vesicle biology, and related subdomains represented in Cluster 16. Nodes represent proteins, and node size reflects network degree (central connectivity). Highly connected (hubs) Proteins like PRKN, APP and MAPT are highlighted.

(C) Boxplots showing PARK7 z-score distributions stratified by sex across (i) all subjects, (ii) AD-only, and (iii) AsymAD-only groups. Although PARK7 is strongly upregulated in Cluster 16 (Fig. 5E), sex-stratified comparisons across the entire dataset reveal no significant differences between males and females (all $P > 0.5$). These results confirm that elevated PARK7 expression is cluster-specific rather than broadly sex-driven.

**Suppl. Fig. 12: Clinical and pathological characteristics of individuals across graph-based clusters.**

(A) Violin plots showing the distribution of six clinical and neuropathological measures—age at death, APOE4 genotype, Braak stage, CERAD score, MMSE score, and postmortem interval (PMI)—across the major clusters identified by graph-based analysis. Each point represents an individual, colored by diagnosis and sex (AD female, AD male, AsymAD female, AsymAD male). The distributions illustrate the heterogeneity of clinical and pathological features across clusters and highlight clusters with distinct demographic or pathological signatures.

(B) Heatmap summarizing pairwise Wilcoxon rank-sum tests between clusters for each clinical and pathological variable. Rows and columns represent all cluster pairs, and cell shading (with asterisks) denotes statistically significant differences after multiple-testing correction ($p < 0.05$), with darker colors indicating smaller p-values. Cluster 16 showed a significant association with age at death, with all individuals in this cluster belonging to the highest age group (≥90 years). Cluster 2, enriched for AsymAD individuals, showed a significant positive association with MMSE scores, consistent with the higher cognitive performance typically observed in this diagnostic category.

**Suppl. Fig. 1.**

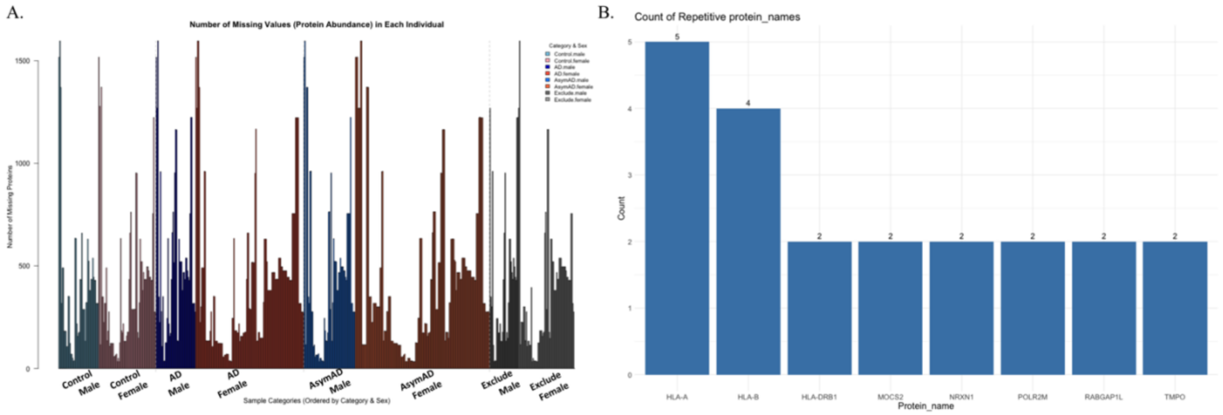


**Suppl. Fig. 2.**

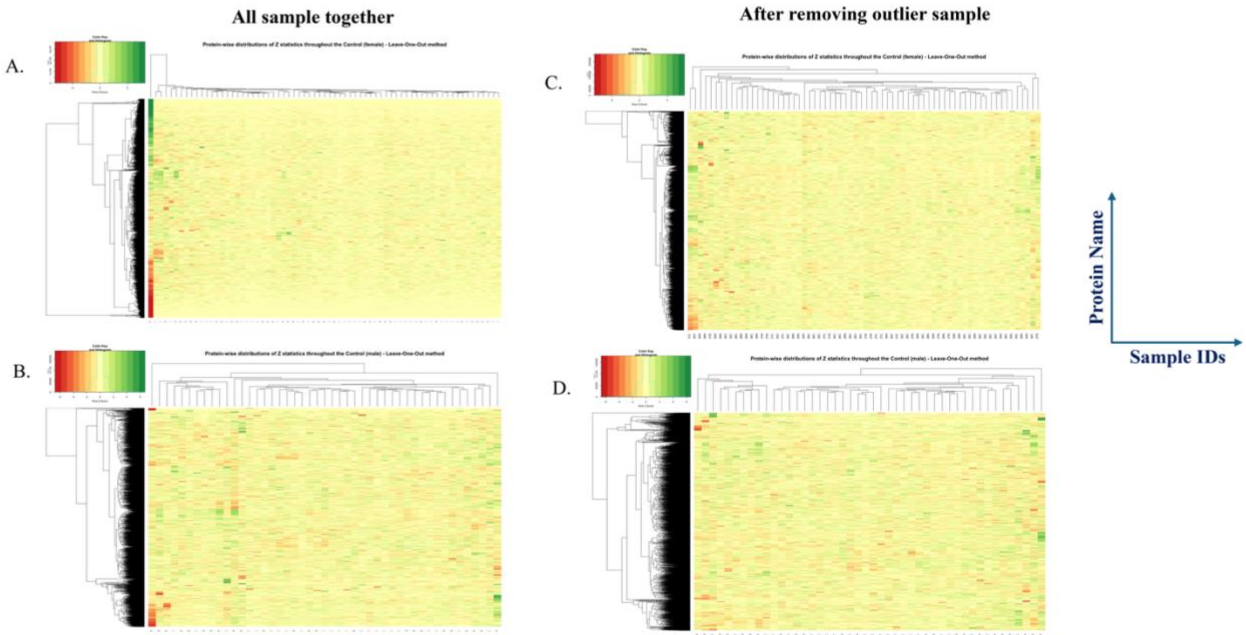

**Suppl. Fig. 3.**

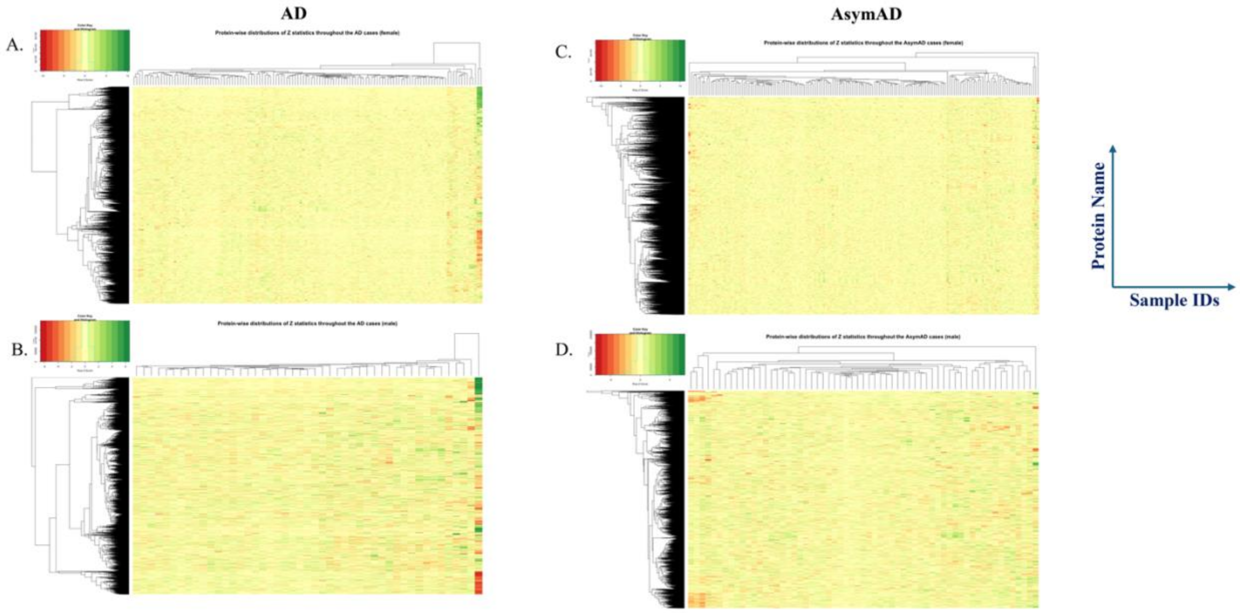


**Suppl. Fig. 4.**

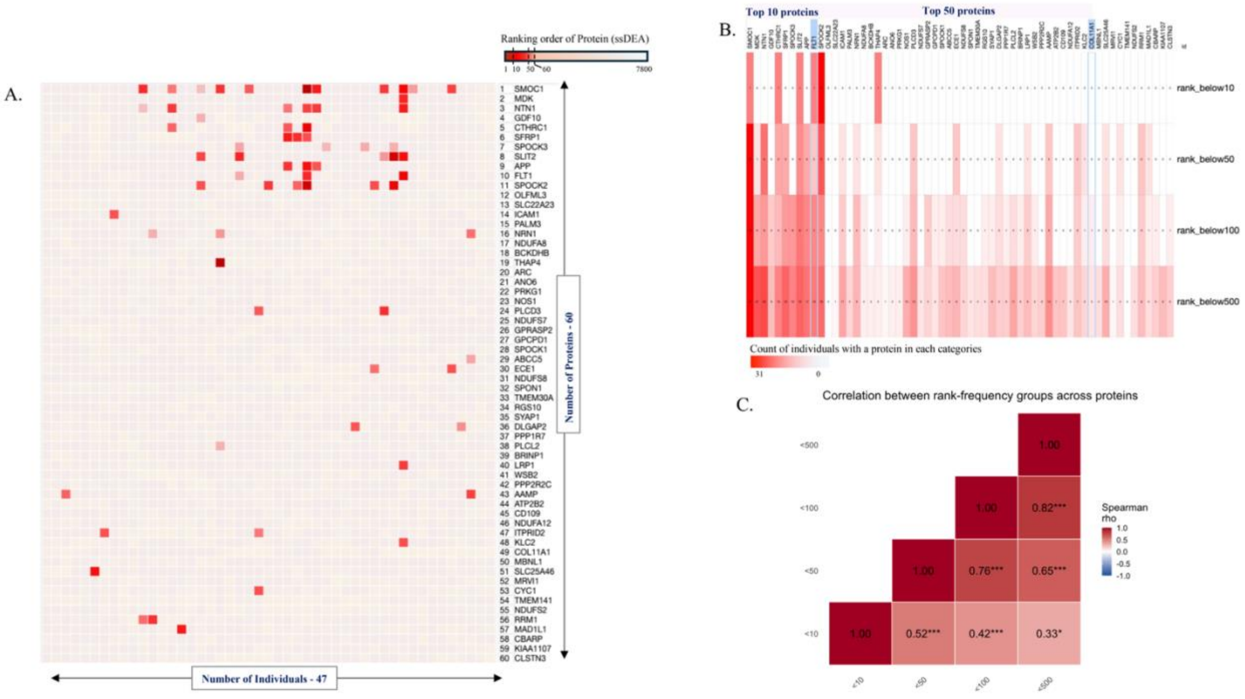

### Suppl. Fig. 5.

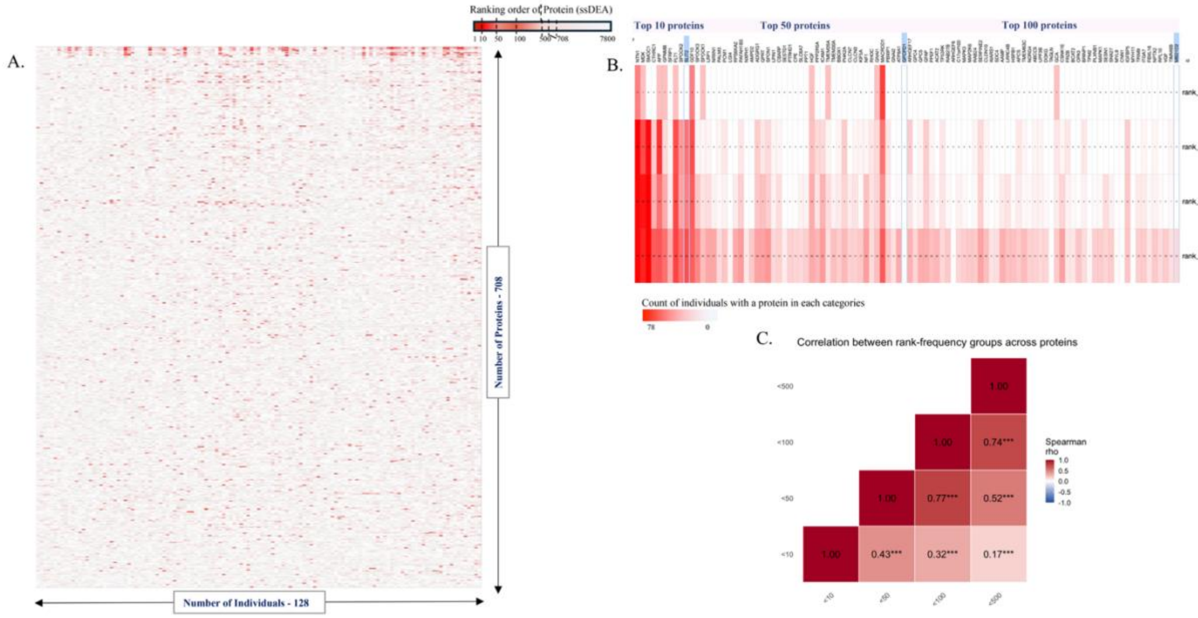


### Suppl. Fig. 6.

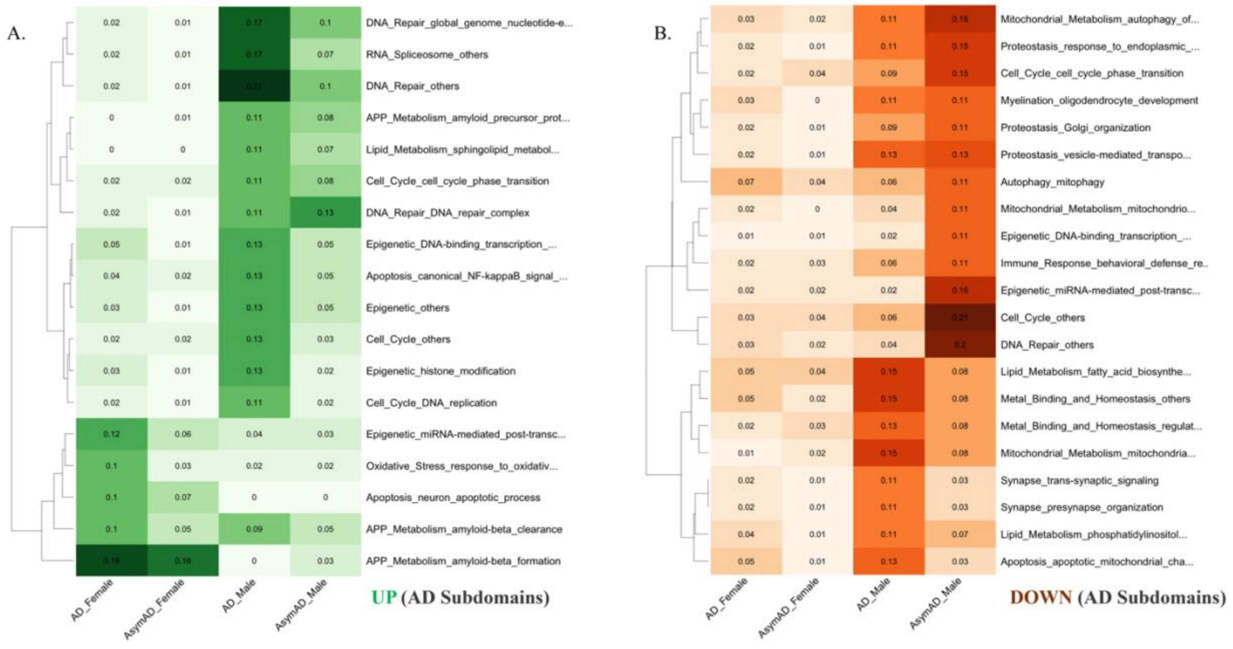

### Suppl. Fig. 7.

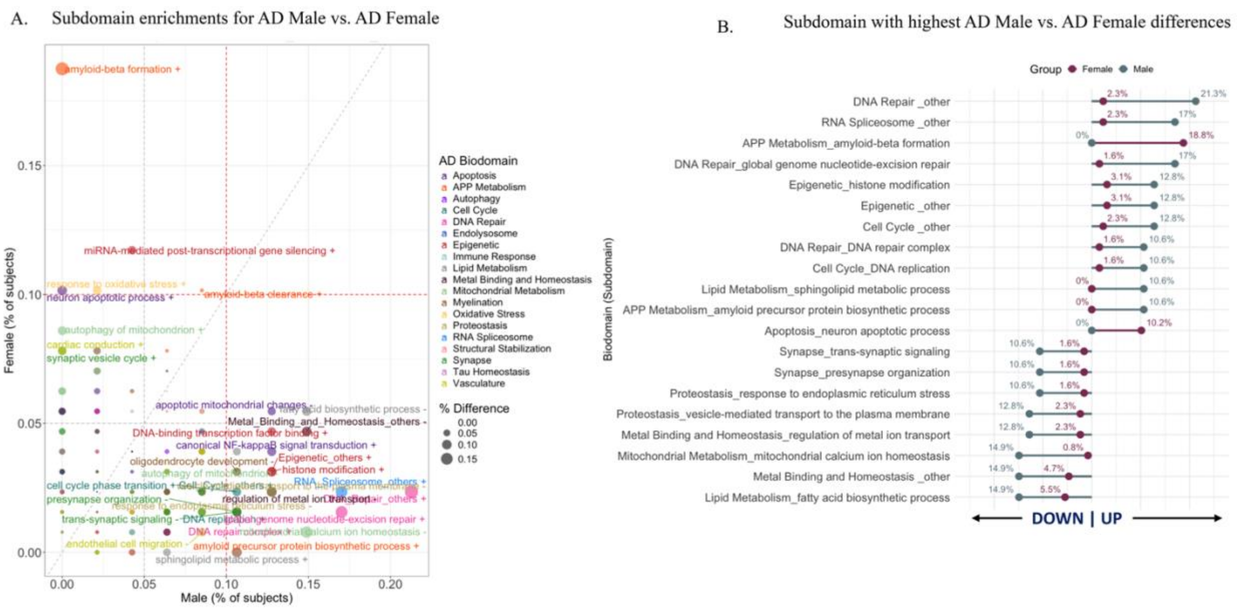


### Suppl. Fig. 8.

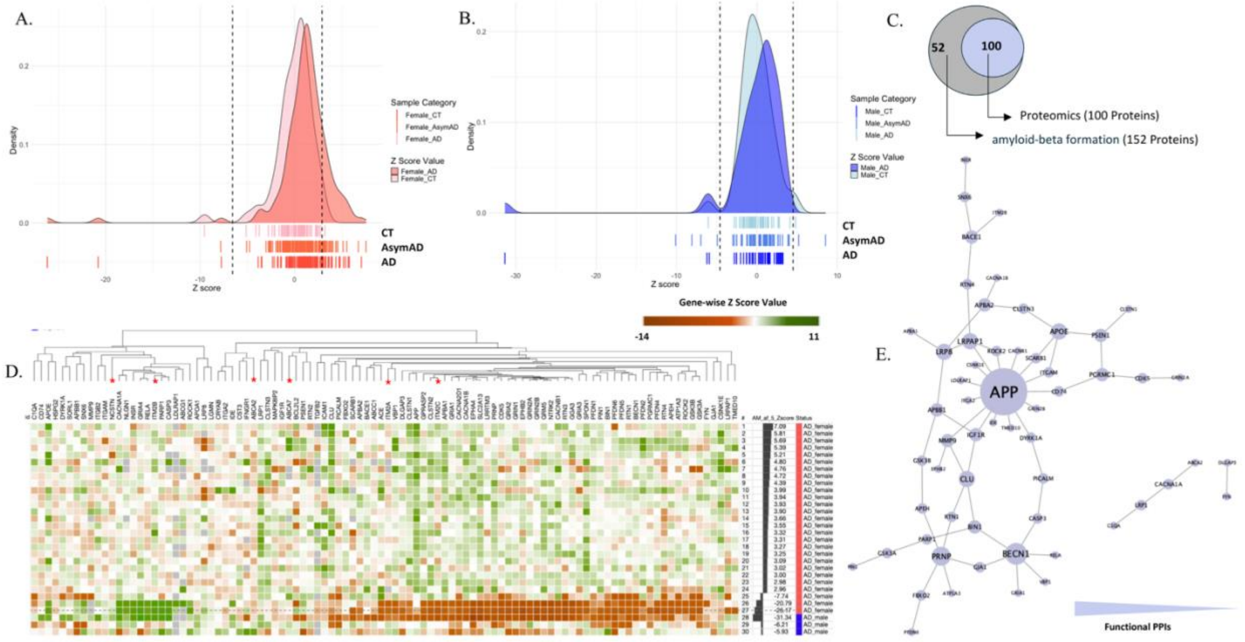

### Suppl. Fig. 9.

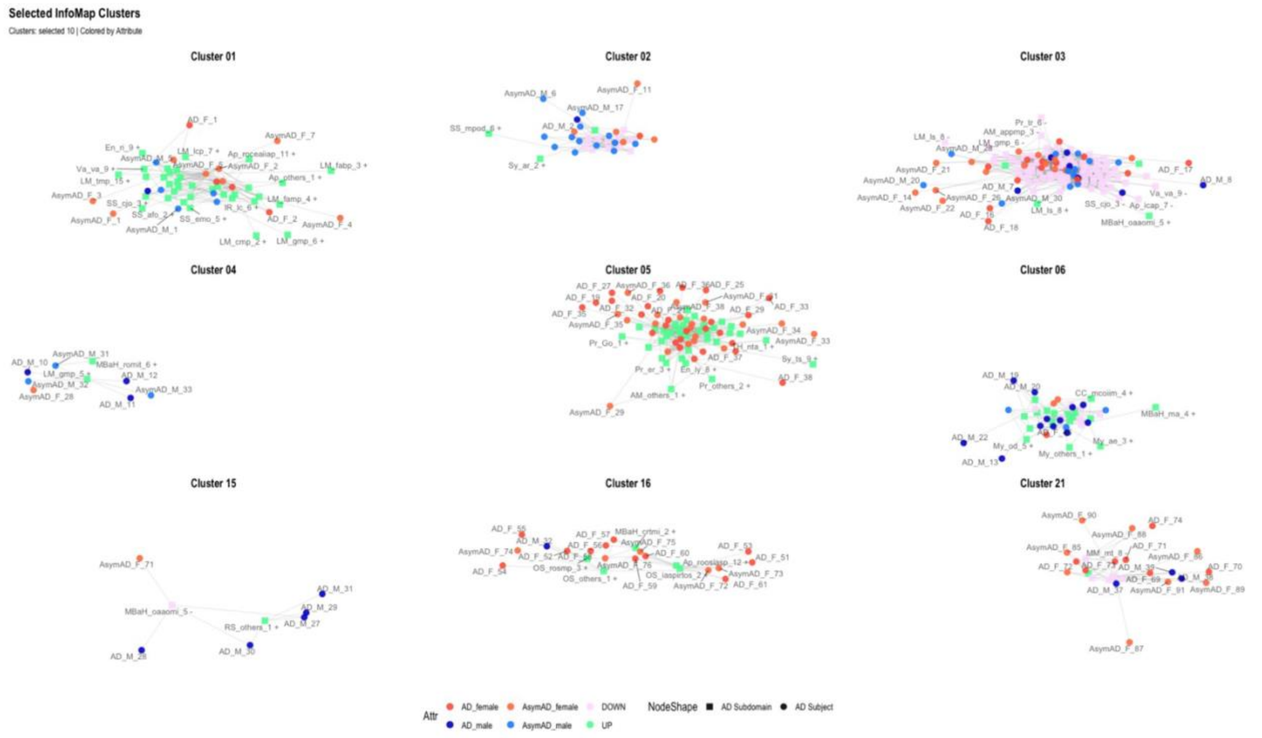


### Suppl. Fig. 10.

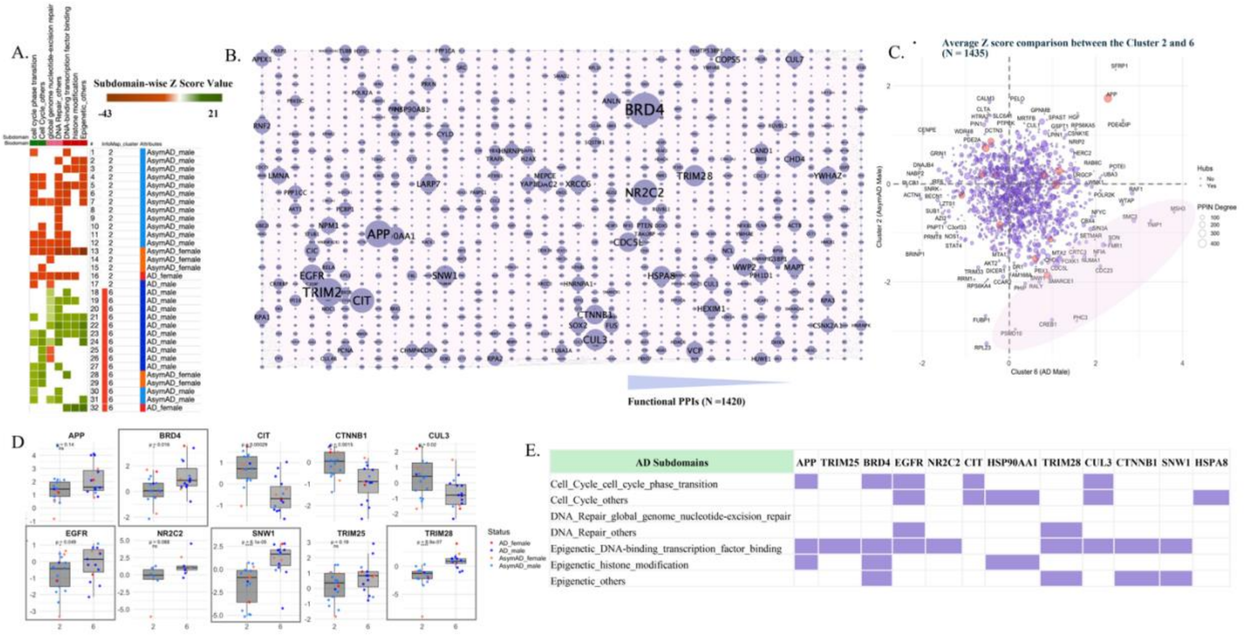

**Suppl. Fig. 11.**

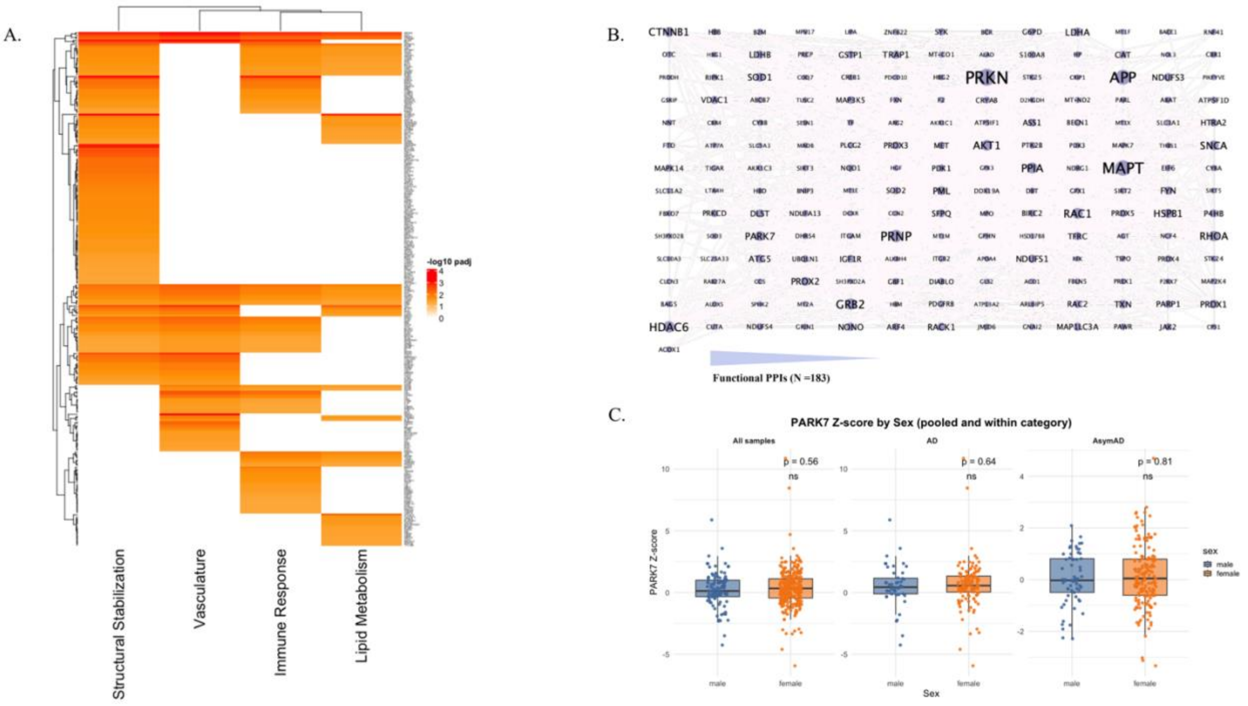


**Suppl. Fig. 12.**

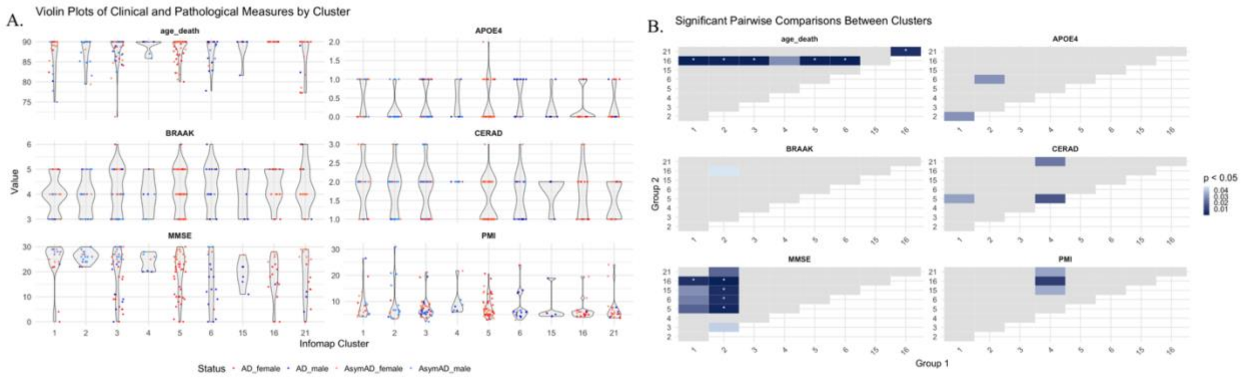
